# A global data-driven census of *Salmonella* small proteins and their potential functions in bacterial virulence

**DOI:** 10.1101/2020.05.26.116038

**Authors:** Elisa Venturini, Sarah L. Svensson, Sandra Maaß, Rick Gelhausen, Florian Eggenhofer, Lei Li, Amy K. Cain, Julian Parkhil, Dörte Becher, Rolf Backofen, Lars Barquist, Cynthia M. Sharma, Alexander J. Westermann, Jörg Vogel

## Abstract

Small proteins are an emerging class of gene products with diverse roles in bacterial physiology. However, a full understanding of their importance has been hampered by insufficient genome annotations and a lack of comprehensive characterization in microbes other than *Escherichia coli*. We have taken an integrative approach to accelerate the discovery of small proteins and their putative virulence-associated functions in *Salmonella* Typhimurium. We merged the annotated small proteome of *Salmonella* with new small proteins predicted with *in silico* and experimental approaches. We then exploited existing and newly generated global datasets that provide information on small open reading frame expression during infection of epithelial cells (dual RNA-seq), contribution to bacterial fitness inside macrophages (TraDIS), and potential engagement in molecular interactions (Grad-seq). This integrative approach suggested a new role for the small protein MgrB beyond its known function in regulating PhoQ. We demonstrate a virulence and motility defect of a *Salmonella* Δ*mgrB* mutant and reveal an effect of MgrB in regulating the *Salmonella* transcriptome and proteome under infection-relevant conditions. Our study highlights the power of interpreting available “omics” datasets with a focus on small proteins, and may serve as a blueprint for a data integration-based survey of small proteins in diverse bacteria.

## INTRODUCTION

Small proteins, loosely defined as shorter than 100 amino acids (aa), are being increasingly implicated in regulating major biological processes in all kingdoms of life (Makarewich and Olson, 2017; Saghatelian and Couso, 2015; Storz et al., 2014). In bacteria, small proteins have long been known to perform both structural and regulatory functions in ribosomal subunits. Similar is the case of small protein members of toxin-antitoxin (TA) systems, particularly type-II, where both the toxin and the antitoxin are proteins (Harms et al., 2018). However, starting with the discovery almost two decades ago of previously overlooked conserved small open reading frames (sORFs) in the *Escherichia coli* chromosome (Wassarman et al., 2001), interest in other potential roles of bacterial small proteins has been increasing (Miravet-Verde et al., 2019; Sberro et al., 2019; Storz et al., 2014). In-depth characterization of individual small proteins has since revealed an unexpected diversity of functions in several different species. For example, the *E. coli* small protein MgtS (31 aa) indirectly increases the intracellular level of magnesium by binding and regulating the activity of the magnesium importer MgtA (Wang et al., 2017) and the cation-phosphate symporter PitA (Yin et al., 2019). SgrT (43 aa), a small protein encoded by the dual-function sRNA SgrS in Enterobacteriaceae, inhibits the activity of the major glucose transporter PtsG under sugar-phosphate stress (Lloyd et al., 2017). In *Listeria monocytogenes*, Prli42 (31 aa) is essential for survival in macrophages as a previously overlooked member of the stressosome (Impens et al., 2017). The variety of functions that have so far been attributed to the few characterized small proteins suggests that much remains to be discovered. Moreover, a clear picture of how many *bona fide*, translated sORFs are encoded even by otherwise well-studied model bacteria is currently lacking.

While the prevalence and functionality of bacterial small proteins remain best understood in the non-pathogenic *E. coli* strain K12, there is increasing evidence for small protein functions in related enteric pathogenic bacteria, especially in *Salmonella enterica* serovar Typhimurium (henceforth, *Salmonella*). Pre-genomic work showed that these two model species of microbiology differ by several large genetic regions that *Salmonella* acquired in its evolution towards becoming an intracellular pathogen of eukaryotic hosts (Groisman and Ochman, 1996). For example, the *<u>S</u>almonella* <u>p</u>athogenicity islands 1 and 2 (SPI-1, SPI-2) each encode a type-III secretion system (T3SS) that translocates its corresponding effector proteins into the host cell where they modulate host cellular processes to the bacterium’s benefit (Jennings et al., 2017; Patel and Galán, 2005). However, early genomic comparisons showed that the large majority of genetic differences between the *E. coli* and *Salmonella* genomes are small (Parkhill et al., 2001), and there are numerous distinctive regions in the *Salmonella* genome that encode different virulence-associated factors (Dos Santos et al., 2019). These loci might harbour previously overlooked small proteins. In other words, a systematic annotation and analysis of small proteins in *Salmonella* promises not only to unveil functions of *E. coli* sORFs that are conserved (or deviate) in a related species, but it might also reveal *Salmonella*-specific small proteins, some of which might contribute to virulence of this pathogen.

Indeed, while only a handful of small proteins have been characterized in *Salmonella*, several of them proved to have a virulence-related function. For example, we recently found CspC and CspE (69 and 70 aa, respectively) to be essential for *Salmonella* pathogenicity and demonstrated a global RNA-binding function for these two small proteins (Michaux et al., 2017). Besides these examples, the small protein MgtR (30 aa) promotes the degradation of MgtC, thereby contributing to titration of the levels of this virulence factor (Alix and Blanc-Potard, 2008). At the same time, together with MgtS (see above), MgtR regulates the activity of the magnesium importer MgtA under Mg^2+^ limiting conditions that *Salmonella* typically experiences inside its host cell (Choi et al., 2012). Also related to infection, PmrD (85 aa) is required for the response of the PmrAB system that confers resistance to antimicrobial peptides (Kox et al., 2000).

The MgrB (47 aa) small protein has been of special interest because it binds and inhibits the PhoP kinase of the PhoPQ two-component system, which is the central regulator of *Salmonella*‘s intracellular virulence program (Lippa and Goulian, 2009). The inhibition of PhoPQ is conserved in *E. coli* where MgrB was shown to localize to the cell membrane, and where this small protein inhibits the kinase activity of PhoQ as part of a negative feedback loop (Salazar et al., 2016), since *mgrB* transcription itself is activated through PhoPQ (Kato et al., 1999). Recently, interest in MgrB has further increased as several reports linked this small protein to antibiotic resistance development in the opportunistic pathogen *Klebsiella pneumoniae* (Kidd et al., 2017; Poirel et al., 2015; Zowawi et al., 2015). These case studies notwithstanding, our knowledge of the role of MgrB in bacterial virulence, and the potential contribution of other small proteins to *Salmonella* infections remains poorly understood.

To close this knowledge gap, we set out to systematically study the virulence-related small proteome of *Salmonella*. We employed both computational predictions (sPepFinder; (Li and Chao, 2020)) and experimental ribosome profiling data to refine the sORF annotation of *Salmonella*. In a second step, given that numerous global datasets exist for this model pathogen, we re-analysed some of these with a focus on our updated small protein annotation (**Fig. 1**). Specifically, we revisited available data that provide information on gene expression during host cell infection (dual RNA-seq; (Westermann et al., 2016)), on *Salmonella* mutant fitness upon uptake by macrophages (<u>Tra</u>nsposon-<u>d</u>irected insertion <u>s</u>equencing [TraDIS]), as well as gradient profiling by mass spectrometry (Grad-seq) data (Smirnov et al., 2016) that indicate a possible involvement of the small proteins in cytosolic complexes. This integrative reanalysis provides a starting point for identifying small proteins with virulence factor potential. We establish a proof-of-principle with the small protein MgrB, for which we report PhoPQ-dependent and possibly PhoPQ-independent functions in *Salmonella* virulence. Altogether, our study exemplifies how the combination of ribosome profiling, computational prediction and the reanalysis of the wealth of existing omics datasets is key in the functional characterization of small proteins.

**Figure 1:**
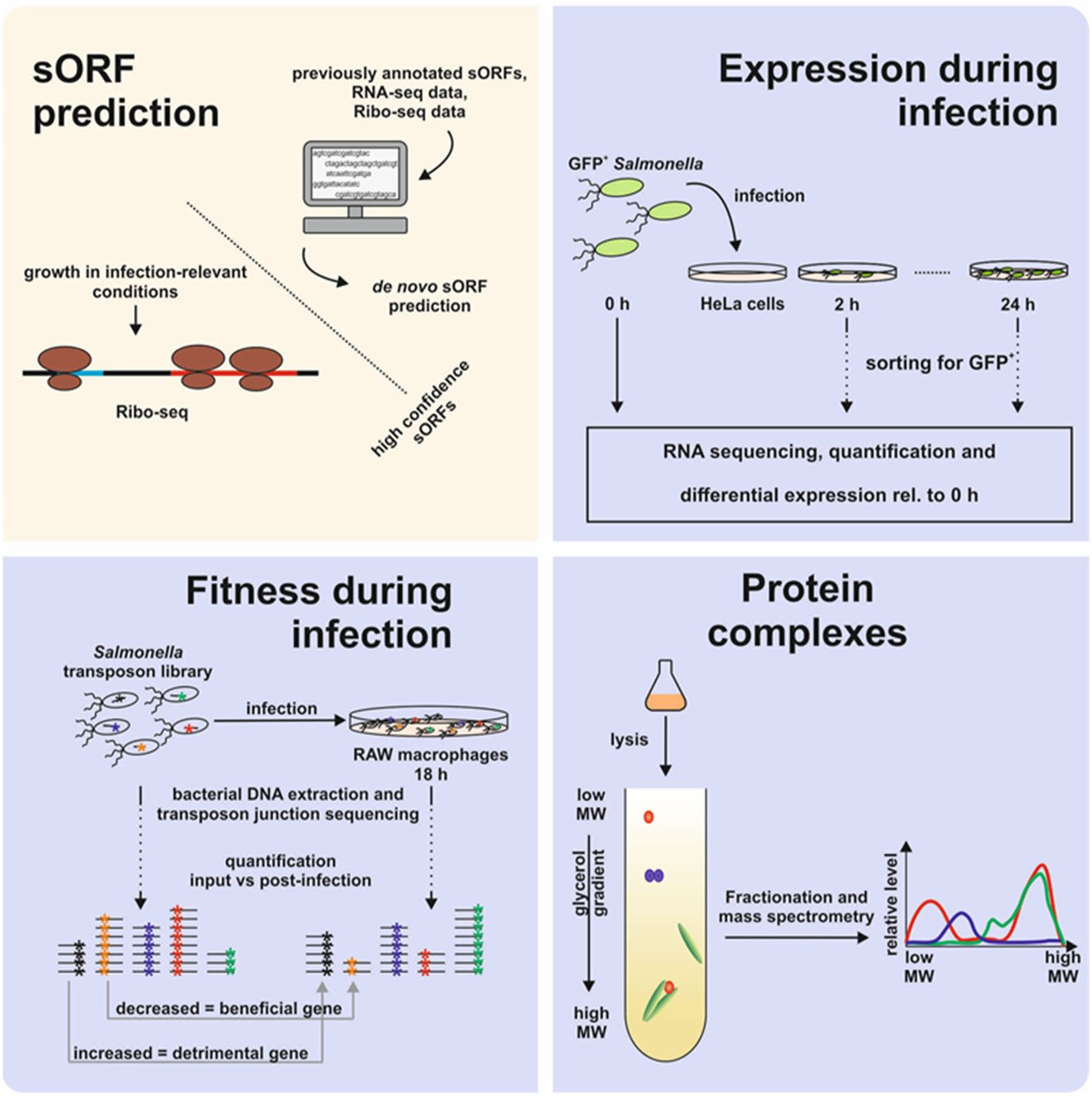
Overview of the in silico and experimental approaches included in the present study. For the prediction of novel small proteins (top left), we included results from sPepFinder, a pipeline for the de novo prediction of sORFs in bacterial genomes, as well as data from ribosome profiling performed on *Salmonella* grown in infection-relevant conditions. To prioritize candidates for functional characterization, we explored data derived from dual RNA-seq, transposon-directed insertion sequencing (TraDIS), and gradient profiling (Grad-seq). Dual RNA-seq (top right) shows the expression pattern of each gene during infection thanks to the enrichment of host cells infected with GFP-expressing *Salmonella*. Total RNA from these cells is sequenced and the bacterial transcriptome is analysed, calculating the abundance of each transcript relative to the bacterial inoculum (0 h). TraDIS (bottom left) informs on sORFs whose disruption by transposon insertion affect fitness during infection. Gradient profiling (or Grad-seq, bottom right) makes use of a linear glycerol gradient to separate the soluble complexes of *Salmonella* lysate based on shape and molecular weight. Mass spectrometry analysis of the gradient fractions shows the sedimentation pattern of each protein, and correlation of the distribution profiles of individual proteins provides information about their potential interactome.

## RESULTS

### Biocomputational search for novel sORF candidates in *Salmonella*

The *Salmonella* SL1344 genome encodes 4657 annotated proteins (as of April 2019), of which 470 are shorter than 100 aa. This suggests that small proteins are underannotated compared to the rest of the genes, a situation similar to that of *E. coli* (**Fig. 2a**). The functional classes with the highest frequency of small proteins in *Salmonella* are small ribosomal proteins (22), T3SS-associated proteins (eight), cold-shock proteins (five), and members of toxin-antitoxin systems (four). All the same, however, functional annotations are missing for the vast majority of *Salmonella* small proteins.

**Figure 2:**
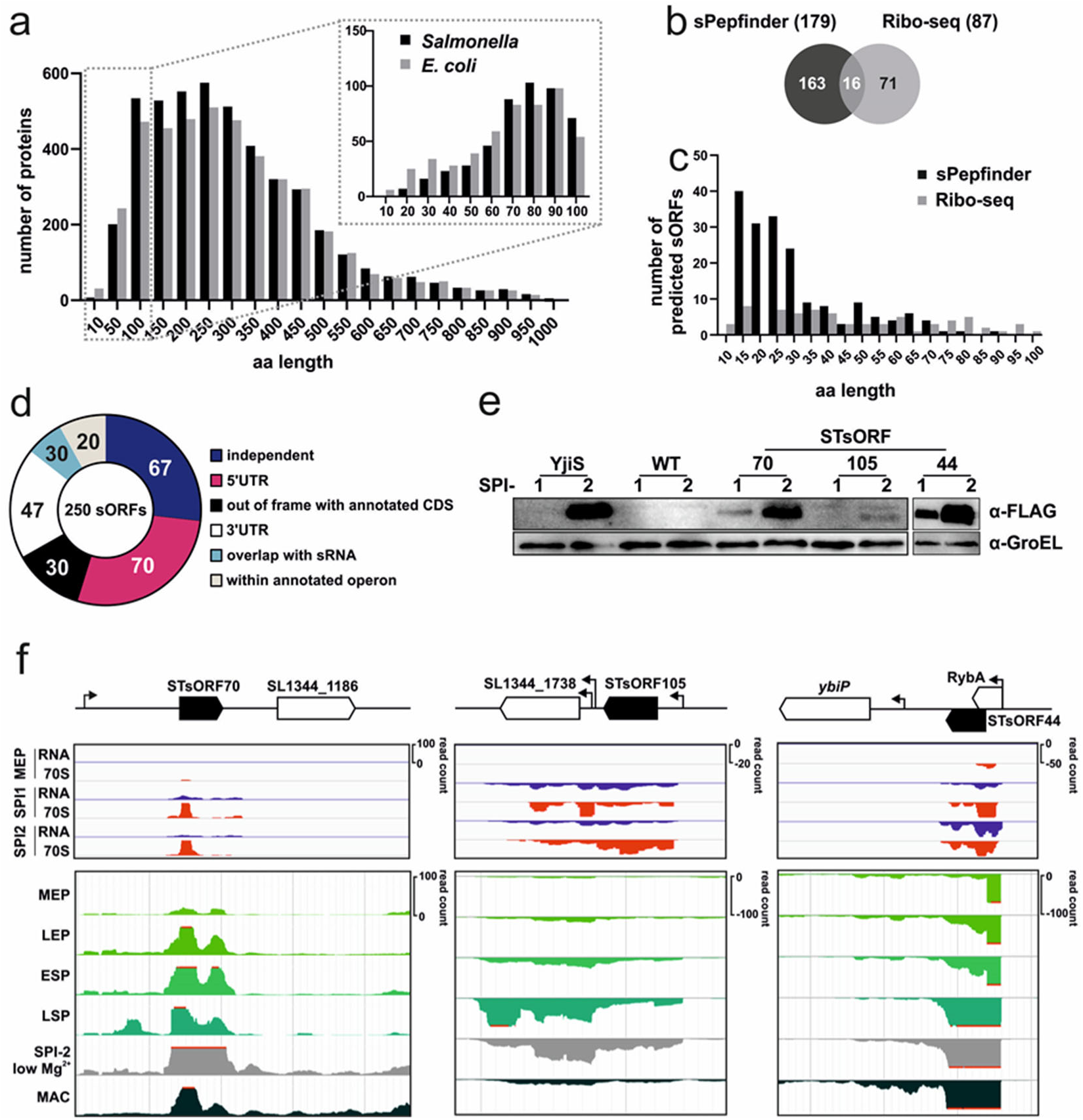
STsORF predictions and validations. **a**, Length distribution of the proteins (up to 1000 aa) annotated in *Salmonella* SL1344 and *E. coli* K12 on UniProt as of April 2019. The upper histogram is a zoom-in of the length distribution of the small proteins annotated in the two organisms. **b**, Overlap of the sORF predictions from sPepFinder and Ribo-seq data. **c**, Length distribution of the new small proteins identified with sPepFinder and Ribo-seq data. The 16 sORFs common to both approaches are grouped with sPepFinder. **d**, Genomic distribution of the new sORFs relative to currently annotated genes. **e**, Validation of three small proteins. STsORF70, STsORF105, and STsORF44 (annotated in NCBI as *mntS*) were chromosomally tagged with the SPA epitope. Each strain was grown in LB to an OD_600_ of 2.0 (SPI-1 condition) or in SPI-2 medium, as indicated by lanes marked with 1 and 2, respectively. The wild-type *Salmonella* strain was included as a background control for the antibody, while the YjiS-SPA tag strain was included as a positive control for a known small protein. **f**, Schematic representation of the validated STsORFs (black arrows) relative to annotated genes (white arrows) and TSSs (thin arrows). Top panels: Expression of the three validated sORFs as detected by Ribo-seq in MEP (mid-exponential phase), SPI-1- and SPI-2-inducing conditions. RNA = total RNA; 70S = ribosome footprints. Bottom panels: expression levels from SalComMac (Srikumar et al., 2015) of the three validated sORFs from **e**. MEP: mid-exponential phase; LEP: late exponential phase; ESP: early stationary phase; LSP: late stationary phase; SPI-2 low Mg^2+^: SPI-2-inducing; MAC: inside macrophages.

In an attempt to bring the sORF annotation in *Salmonella* closer to completion, we searched for previously overlooked small protein candidates by combining computational sORF predictions with experimental data. sPepFinder is a machine learning-based approach for the identification of bacterial sORFs (Li and Chao, 2020). Briefly, sPepFinder uses a support vector machine and 29 features, including a thermodynamic model of ribosome binding sites and the frequency of hydrophobic amino acids, extracted from a training set of annotated bacterial small proteins from ten species. We applied sPepFinder to the *Salmonella* strain SL1344 genome with a length cut-off of 100 codons.

After training on the small proteins annotated in *Salmonella*, as well as in other bacterial genomes, sPepFinder predicted further 340 sORFs (**Table S1**). We filtered these to exclude the ones that did not pass the statistical filtering cut-off (SVM probability > 0.9). Furthermore, since sPepFinder predictions are based solely on genomic features, we sought to obtain independent evidence that the identified sORF candidates are indeed expressed. To this end, we interrogated the SalComMac database (Srikumar et al., 2015), which contains transcriptomic data of *Salmonella* grown in diverse conditions, including those related to pathogenesis (e.g. during growth inside macrophages). Overall, we identified 179 potential new sORFs. For our newly predicted genes, we propose the nomenclature ‘STsORF’ followed by sequential numbers, based on the position on the genome.

Some examples of condition-specific expression patterns include the candidate STsORF145 (31 aa), expressed during *Salmonella* growth in different shock conditions (**Fig. S1a**). A substantial number of sORF candidates were expressed under infection-related conditions, namely when *Salmonella* was grown under an invasion genes-inducing condition (referred to as “SPI-1” condition or late exponential phase; exemplified by STsORF55, 34 aa), or in minimal medium mimicking the intravacuolar environment (“SPI-2 low Mg^2+^”; e.g. STsORF72, 35 aa) (**Fig. S1b**). In total, based on the combination of *in silico* genome-based predictions with available transcriptomic datasets, we added 179 novel candidates with strong evidence for expression to the annotated *Salmonella* sORFs.

### Experimental prediction of *Salmonella* sORFs

To validate the sPepFinder candidates and discover additional sORFs based on association of their mRNAs with translating ribosomes, we used the genome-wide experimental approach of ribosome profiling by sequencing (Ribo-seq; (Ingolia et al., 2012; Ingolia et al., 2009)). By sequencing the ∼30 nt mRNA fragments that are protected from nuclease digestion by actively translating ribosomes, Ribo-seq provides a global picture of the transcripts being translated at a given time, enabling the determination of ORF boundaries, and has already successfully been applied for sORFs annotation (Canestrari et al., 2020; Weaver et al., 2019). Ribo-seq thus overcomes limits of traditional mass spectrometry approaches inherent in their dependence on predicted amino acid sequences, and circumvents the need for epitope-tagging or antibody production for Western blotting of small proteins, which are both time consuming and technically challenging (Makarewich and Olson, 2017).

We applied Ribo-seq to *Salmonella* grown *in vitro* under three conditions (mid-exponential phase in LB (MEP), SPI-1-inducing, or SPI-2-inducing). The resulting data were analysed with REPARATION (Ndah et al., 2017), a tool for bacterial gene annotation based on Ribo-seq data. After filtering for sORF predictions with a translational efficiency (TE) > 0.5 in at least one of the three conditions and visual inspection of the sequencing coverage, computational prediction by REPARATION resulted in 87 novel, and 224 known, sORFs with a size between 10 and 99 aa.

The overlap between sORF candidates predicted by Ribo-seq and sPepFinder was small (16 sORFs, **Fig. 2b**), likely due to the fact that both approaches have different biases. Since sPepFinder is trained on properties of known small proteins, its predictions could be improved by more comprehensive annotations. Similarly, Ribo-seq performed on cells grown in multiple conditions could also result in more candidates, and several false-positives arise from transcripts associated with the ribosome without being translated. Nevertheless, to capture as many sORF candidates as possible, we decided to include not only the sixteen sORF candidates called by both approaches, but also the 163 and 71 candidates called exclusively by either sPepFinder or Ribo-seq, respectively. As these candidates passed the statistical cut-offs applied for the respective datasets, only subsequent thorough validation will inform on the rate of false-positive occurrences. The size distribution of the resulting 250 novel sORF candidates is shown in **Fig. 2c**. Inspection of transcription start site (TSS) annotations (Kröger et al., 2013) revealed a high prevalence of 5’UTR-encoded sORFs, as well as independent sORFs (**Fig. 2d**).

### Validation of new sORF candidates

We next set out to validate the translation of several representatives of the combined set of 250 novel sORF candidates. Three of the newly predicted candidates were chromosomally tagged with the sequential peptide affinity (SPA) tag (Zeghouf et al., 2004) – an epitope previously used to detect small bacterial proteins by immunoblotting (Baek et al., 2017; Weaver et al., 2019). The C-terminal fusion strains were grown under SPI-1- or SPI-2-inducing conditions, and total protein samples were harvested and analysed by Western blot. We included the small protein YjiS (54 aa) as a positive control, a validated small protein known to be induced under SPI-2 conditions (Baek et al., 2017). In this way, we confirmed the translation of the 5’UTR-encoded STsORF70, the intergenically-encoded STsORF105, and STsORF44 (a.k.a. MntS; (Baek et al., 2017)), which overlaps with the sRNA RybA (**Fig. 2e**). All three tagged STsORFs were predicted from Ribo-seq data, with STsORF44 also being called by sPepFinder. The CDSs of the three sORFs are highly conserved within *Salmonellae* (**Fig. S2**), further supporting a function in the cell. Moreover, all three were induced in the SPI-2 compared to the SPI-1 condition, in line with the expression of their mRNAs (**Fig. 2f**), and providing evidence for their potential involvement in *Salmonella* virulence. For further analysis, we merged the 250 newly identified small proteins with the 470 annotated *Salmonella* sORFs, summing up to a refined small proteome featuring 720 entries.

### sORF expression kinetics during host cell infection

Induction of sORF transcription during infection may be an indication of a virulence-related function of the corresponding protein. Multiple novel and previously-annotated sORFs are highly expressed under infection-related conditions, both in SalComMac (Srikumar et al., 2015) and in our Ribo-seq data. To further pinpoint small proteins with a likely role in virulence, we re-analysed global gene expression in intracellular *Salmonella* during epithelial cell infection (Westermann et al., 2016) with our updated sORF annotation. This revealed that 280 annotated sORFs were differentially expressed (FDR < 0.05) compared to their expression levels in the inoculum (**Fig. 3a**, **Table S2**). The three most highly induced known sORFs at this early infection stage encode members of the T3SS apparatus (SsaS and SsaI; 88 and 82 aa, respectively) and the uncharacterized YjiS protein (see above). Additionally, expression of *mgrB* (a.k.a. *yobG* in *Salmonella* (Lippa and Goulian, 2009)) peaked at 2 h p.i. (log_2_FC = 3 compared to pre-infection conditions), but remained elevated up to 16 h p.i.

**Figure 3:**
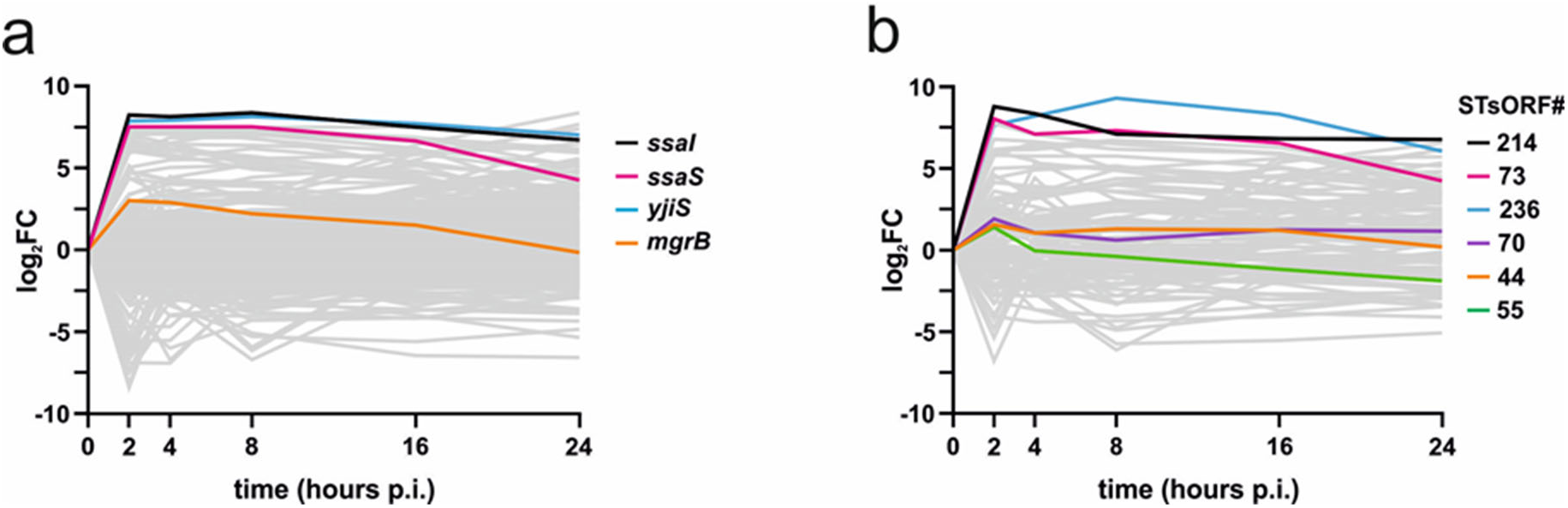
Small protein expression patterns during infection. **a**, Expression of 280 annotated sORFs detected in dual RNA-seq and significantly regulated (FDR < 0.05) in comparison to the inoculum (0 h). The highlighted transcripts are the top three most highly-induced sORFs (log_2_FC > 7 at 2 h p.i.), as well as *mgrB* (log_2_FC = 3 at 2 h). **b**, Expression of 109 new sORFs detected as significantly regulated with respect to the inoculum (FDR < 0.05) over the course of infection. The highlighted transcripts are the top three most highly-induced STsORFs (STsORF214, STsORF73 and STsORF236), the two validated STsORF70 and STsORF44, and the SPI-1-induced STsORF55.

Among the 109 differentially expressed new STsORFs, three (STsORF214, STsORF73, and STsORF236) were highly induced during infection (**Fig. 3b**). STsORF214 (16 aa), encoded in the 5’UTR of *mgtC* (**Fig. S3a**), is expressed with a similar pattern to its downstream gene, a known *Salmonella* virulence factor (Lee and Lee, 2015). Further analysis indicated that STsORF214 is a homolog of the *Salmonella* 14028s protein MgtP, one of the two small proteins encoded in the 5’UTR of *mgtC* that regulated its transcription (Lee and Groisman, 2012). MgtC is missing from the *Salmonella* SL1344 annotations, and was thus included in our predictions. STsORF73 (13 aa) is encoded within the sRNA STnc1480 (**Fig. S3b**), whose expression was shown to be PhoP- and SlyA-dependent (Colgan et al., 2016). STsORF236 (32 aa) is encoded downstream of the acid phosphatase gene *phoN* and is possibly expressed from an annotated TSS internal to *phoN* (**Fig. S3c**). Two of the novel sORFs that we validated as induced under SPI-2 conditions by Western blotting (**Fig. 2e**) — STsORF70 (41 aa) and STsORF44 (42 aa) — were also mildly upregulated during infection (**Fig. 3b**). Small proteins whose mRNAs were downregulated during infection included STsORF55, a 34-aa protein encoded within the 3’UTR of *pipC* (**Fig. S1b**). Its strong repression at 24 h p.i. (log_2_FC = −1.87) reflects the expression pattern of *pipC* itself, encoding the chaperone of the SopB effector protein required for host cell invasion (Darwin et al., 2001).

### Infection phenotypes of sORF disruption mutants

To further narrow in on sORFs with potential functions in virulence, we generated TraDIS data from *Salmonella* infection of macrophages to identify genes whose disruption affected *Salmonella* fitness during infection. To this end, the composition of a transposon mutant pool 20 h after uptake by murine RAW.B macrophages was compared to its composition in the inoculum (**Table S2**). For each gene targeted by transposon insertion, a positive fold-change indicates that the given mutant was over-represented in the pool after infection compared to the input, suggesting the loss of the protein to be beneficial for virulence. A negative fold-change instead indicates that the corresponding protein might be required for infection. As expected, mutants of the *rfa/rfb* clusters, involved in lipopolysaccharide (LPS) O-antigen assembly, were strongly enriched after infection (log_2_FC up to 5.8), in line with previous findings (Zenk et al., 2009). Conversely, mutants of SPI-2 genes were depleted from the pool (e.g. *ssaV* and *sseC*, both with a log_2_FC = −1.2), in accordance with their known requirement for intracellular survival (Browne et al., 2008; Klein and Jones, 2001).

Even though random mutagenesis is inherently biased against small ORFs, our transposon library included mutants for 517 of the 720 sORFs. Seven sORFs passed the applied filtering criteria for an infection phenotype (q-value < 0.05 and |log_2_FC| > 1, **Fig. 4**, **Table S2**). Transposon disruption of *sseA* (encoding a structural protein of the SPI-2 T3SS), *himD* (encoding the β subunit of integration host factor, IHF), *rpoZ* (a subunit of the RNA polymerase; RNAP), *repY* (regulator of the pCol1B9 plasmid replication initiation protein RepZ), and *mgrB* (also induced during HeLa cell infection, **Fig. 3a**) attenuated infection. In contrast, disruption of *yjiS*, which is one of the most highly induced sORFs in *Salmonella* inside epithelial cells (**Fig. 3a**), led to a hyper-virulent phenotype. Similarly, mutants with transposon insertions in *dcoC* (annotated as a probable 81 aa long oxaloacetate decarboxylase) were enriched after infection.

**Figure 4:**
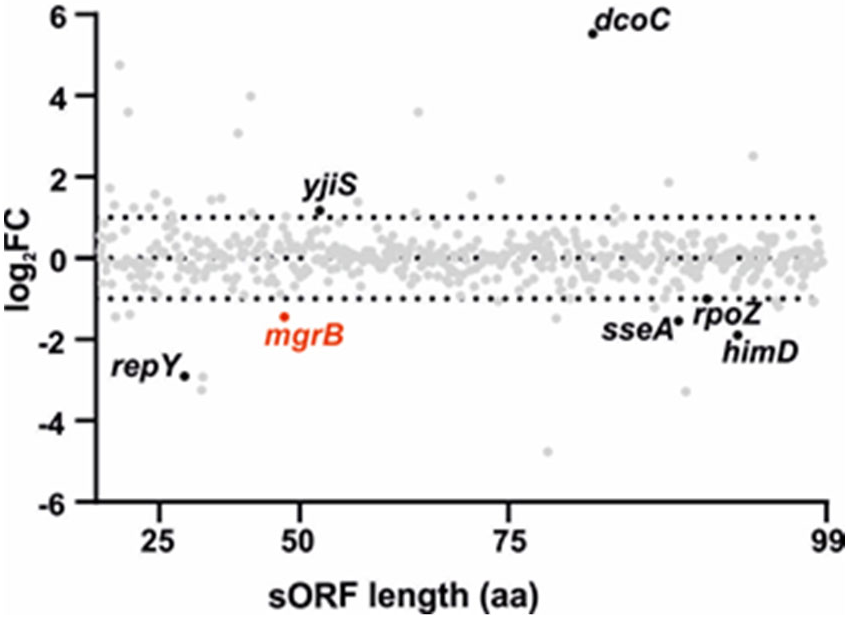
Small protein requirements during infection. Plot of the 517 sORFs shorter than 100 aa with transposon insertions detected in the TraDIS pool. The y-axis displays the log_2_FC of each mutant after quantification in the starting pool vs after 20 h p.i. of RAW.B macrophages (data were collected from two biological replicates). The genes are distributed along the x-axis based on the length (aa). The seven sORFs with |log_2_FC| > 1 and q-value < 0.05 are highlighted.

Loss of RpoZ (91 aa) is likely to affect RNAP function and hence bacterial fitness in general, and interference with the SPI-2 secretion apparatus (SseA, 89 aa) is expected to compromise intracellular survival. Similar is the case of HimD (94 aa), as a fully assembled IHF is required for efficient expression of virulence proteins (Mangan et al., 2006) and for *Salmonella* survival at late stages of macrophage infection (Yoon et al., 2009). In contrast, the virulence phenotype of the uncharacterized sORF *yjiS*, particularly in combination with its intracellular induction, implicates this small protein as a novel factor involved in *Salmonella* virulence.

### Small proteins engaged in larger cellular complexes

Several bacterial small proteins have been found to be integral parts of protein complexes in both the cytosol and the membrane (Storz et al., 2014). To systematically identify molecular interaction partners of *Salmonella* small proteins, we turned to another dataset that provides a global overview of intracellular (ribonucleo-)protein complexes. The Grad-seq approach relies on the separation of soluble cellular complexes on a linear glycerol gradient according to their size and shape, followed by parallel mass spectrometry of each fraction (Smirnov et al., 2016). This provides insights into the cellular complexes a given protein might be engaged in. A small protein not interacting with any other cellular macromolecule would be expected to localize to the low-density fractions of the gradient. In contrast, the localization of a protein in a higher density fraction is indicative of this protein being part of a larger macromolecular complex in the bacterial cell. Of note, while dual RNA-seq and Ribo-seq provide evidence for transcription and translation of a given sORF under a given condition, and TraDIS provides genome-wide genetic evidence for a role during infection, none of these approaches operates at the protein level. In contrast, Grad-seq is coupled to mass spectrometry, and thus allows the detection of the actual gene products of sORFs.

In total, 170 of our 720 small proteins were detected in the Grad-seq dataset from *Salmonella* grown in SPI-1-inducing conditions (**Table S2**). These include all 22 annotated ribosomal proteins, three RNA-binding proteins (CsrA, CspC and CspE), as well as 91 uncharacterized proteins, providing direct evidence for their translation. None of our novel STsORFs were detected in this dataset. As a proof of principle for the validity of Grad-seq-derived information on small proteins, the levels of the small RNAP subunit RpoZ (∼10 kDa, 91 aa) peaked in fraction six, the same fraction where the ∼450-kDa RNAP holoenzyme (RpoB-D; **Fig. 5**) migrates.

**Figure 5:**
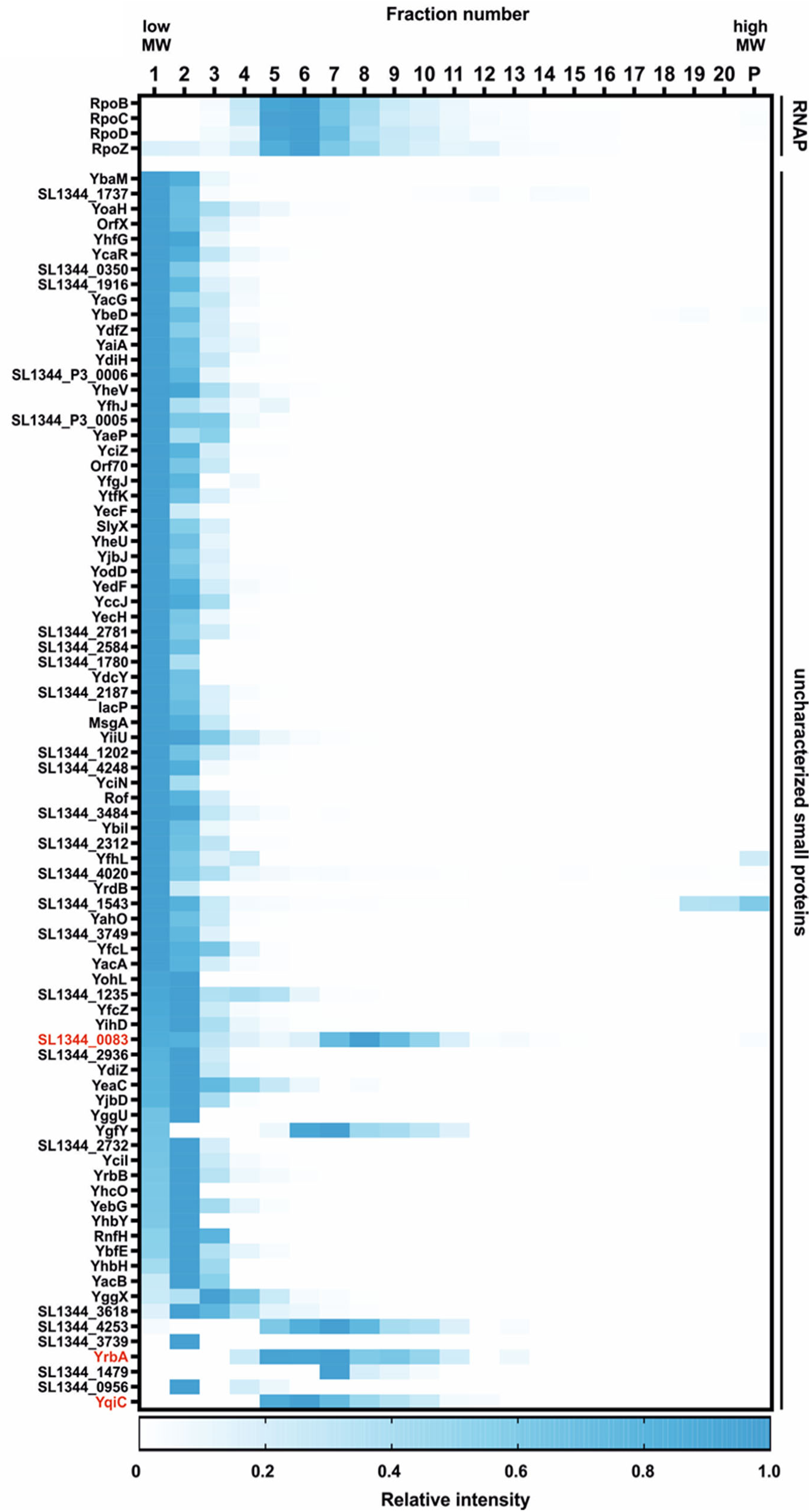
Sedimentation profiles of small proteins. Heat map showing the sedimentation profile of proteins part of the RNA polymerase (RNAP) and of the 82 uncharacterized small proteins detected in Grad-seq under SPI-1-inducing conditions. The intensity in each fraction, including the pellet (“P”), was normalized relative to the fraction with the highest intensity for each protein. The proteins in red are mentioned throughout the text.

Among the detected 89 uncharacterized small proteins, seven were only present in the first fraction, suggesting that they are not engaged in stable molecular interactions in the given experimental condition. The sedimentation profiles of the remaining 82 uncharacterized proteins (**Fig. 5**) showed a variety of patterns. For example, YqiC and YrbA both peaked around fraction six, where the RNAP also sediments. Two peaks were detected for the hypothetical protein SL1344_0083 (one in the low molecular weight fractions and one partially overlapping that of RNAP), suggesting that only a fraction of the cellular proteins are engaged in stable interactions under the analysed condition. The virulence-related small proteins MgrB and YjiS were absent from the Grad-seq dataset, probably for different reasons. While YjiS is not expressed in the growth condition used for Grad-seq (**Fig. 2e**), MgrB is an inner membrane-associated protein and the lysis approach used here does not efficiently recover hydrophobic proteins.

In summary, careful analysis of the above global datasets with focus on our refined sORF annotation pinpointed novel infection-relevant small protein candidates (**Fig. S4**). These include both previously annotated proteins such as YjiS and MgrB, as well as several novel sORFs (STsORF70, STsORF73, STsORF105, and STsORF236).

### *Salmonella* MgrB contributes to macrophage and epithelial cell infection

MgrB was among the small proteins whose mRNA was induced during infection of epithelial cells (dual RNA-seq; log_2_FC = 3 at 2 h p.i.) and whose inactivation attenuated *Salmonella* infection in murine macrophages (TraDIS; log_2_FC = −1.44). Independent infection assays with a clean *mgrB* deletion strain (Δ*mgrB*) supported the hypothesis that this small protein contributes to *Salmonella* virulence not only in macrophages (**Fig. 6a**), but also in epithelial cells (**Fig. S5a**). Particularly, deletion of *mgrB* interfered with the ability of *Salmonella* to enter (**Fig. 6a**, 1 h time point; i.e. before intracellular replication occurs (Steele-Mortimer, 2008)) and to replicate inside both host cell types (**Fig. 6b**, **Fig. S5b**). The virulence defects were (over-)complemented upon transformation of a medium-copy plasmid encoding *mgrB* and its native promoter (*mgrB*^+^ strain; **Fig. 6a**). Of note, the absence of MgrB did not affect *in vitro* growth in LB or SPI-2 medium (**Fig. 6c**), arguing for infection-specific effects rather than a general impact on bacterial fitness. Taken together, these data confirm that the small protein MgrB contributes to *Salmonella* infection of two frequently used cell culture models.

**Figure 6:**
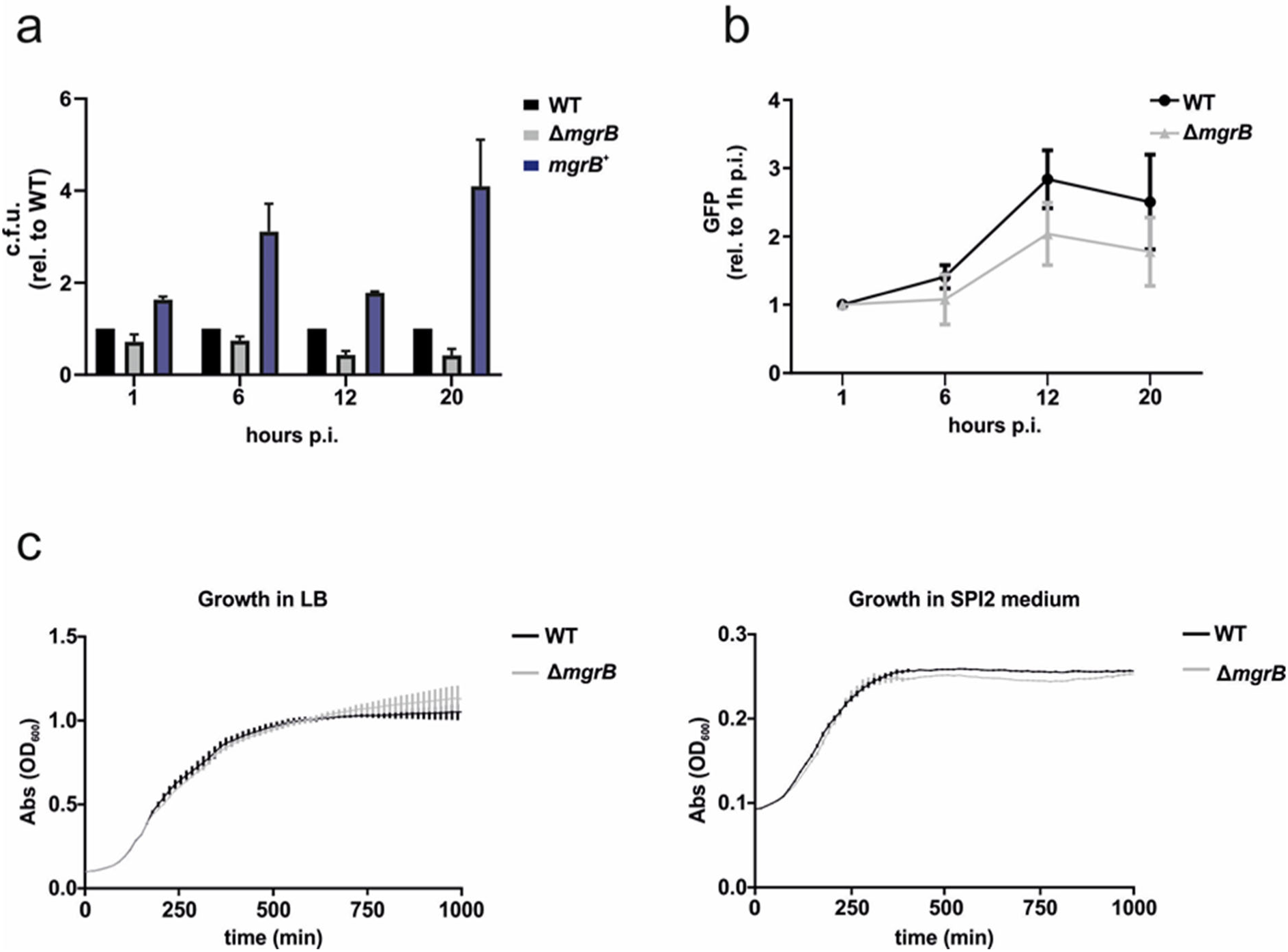
The requirement of MgrB for infection. **a**, Gentamicin protection assay of *Salmonella* wild-type, Δ*mgrB*, or *mgrB^+^* infecting RAW.B macrophages. C.f.u. counts at each time point were normalized to the wild-type strain. **b,** Intracellular replication in RAW.B macrophages of *Salmonella* wild-type or Δ*mgrB* expressing GFP. Fluorescence was normalized to the 1 h time point. For **a** and **b**, data were collected from three biological replicates with error bars indicating standard deviations from the mean. **c**, Growth curve of *Salmonella* wild-type and Δ*mgrB* in LB and SPI-2 medium. For both graphs, data were collected from three biological replicates and bars represent standard deviations from the mean.

### MgrB positively affects the expression of flagella and motility genes

To identify the molecular features that underlie the effect of MgrB on *Salmonella* virulence, we compared the transcriptomes of *Salmonella* wild-type and the Δ*mgrB* strain grown in SPI-2-inducing medium, a condition where MgrB is highly expressed and translated (**Fig. 7a**). Our RNA-seq analysis showed that a subset of genes was downregulated in the Δ*mgrB* mutant relative to the wild-type strain (12 genes with log_2_FC < −2, FDR < 0.05, **Table S3**). These included genes encoding motility- and chemotaxis-related proteins, such as *fliC*, *flgB*, *motB*, *cheR*, and *tar* (**Fig. 7b**).

**Figure 7:**
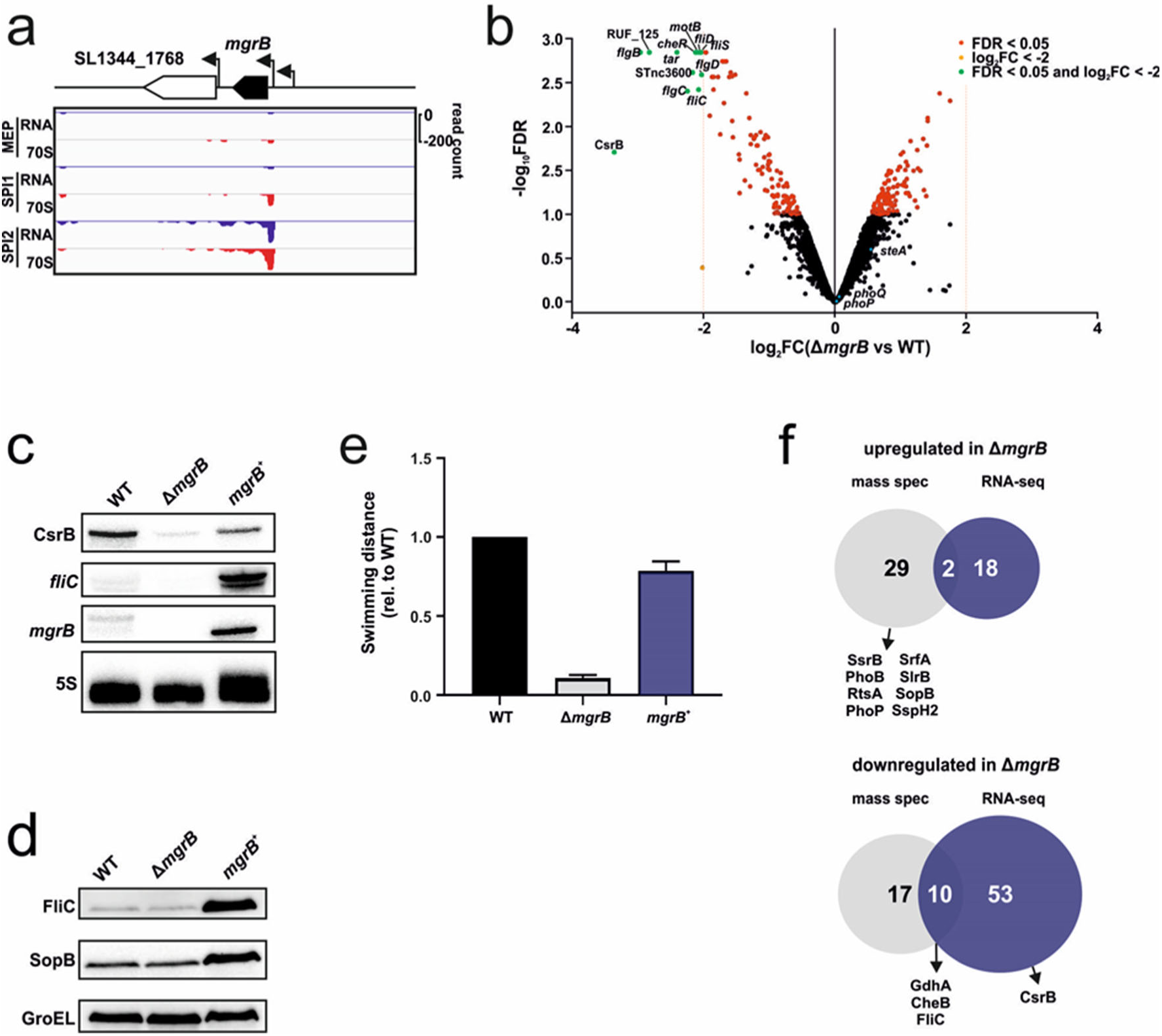
Impact of MgrB on the *Salmonella* transcriptome and proteome. **a**, Expression of *mgrB* as detected by Ribo-seq in MEP (mid-exponential phase), SPI-1-, and SPI-2-inducing conditions. RNA = total RNA; 70S = ribosome footprints. **b,** Comparison of the transcriptomes of *Salmonella* wild-type vs Δ*mgrB* grown in SPI-2 medium. The transcripts with a significant (FDR < 0.05) dysregulation are coloured in red, while those with a |log_2_FC| > 2 are coloured in orange. RNAs that are dysregulated above the threshold of |log_2_FC| > 2 and significant are highlighted in green. Data are from two biological replicates. **c**, Validation via Northern blot of selected transcripts identified as dysregulated by RNA-seq. For this, *Salmonella* wild-type, Δ*mgrB* and *mgrB^+^* were grown in SPI-2-inducing conditions. The figure is representative of three biological replicates. The 5S rRNA was probed for as a loading control. **d**, Validation of the dysregulated proteins FliC and SopB as detected by mass spectrometry in *Salmonella* grown in SPI-2 conditions via Western blot. GroEL was included as a loading control. Strains and replicates are as for panel **c**. **e**, Quantification of swimming distance from the center of Δ*mgrB* and *mgrB^+^* relative to wild-type in SPI-2 medium. Data were generated from three biological replicates and the error bars represent standard deviations from the mean. **f**, Venn diagrams displaying the overlap between upregulated (top) and downregulated (bottom) proteins or transcripts generated from mass spectrometry and RNA-seq analysis in the presence or absence of *mgrB*.

We independently validated the downregulation of *fliC* in Δ*mgrB* compared to wild-type *Salmonella*, both on the mRNA (**Fig. 7c**) and the protein level (**Fig. 7d**) by Northern or Western blot analysis, respectively. Reduced expression of motility genes in the absence of MgrB lead us to hypothesize that the Δ*mgrB* mutant might have a motility defect. Indeed, the Δ*mgrB* mutant was defective in swimming on SPI-2 medium/0.3% agar plates, an effect that could be complemented in *trans* (**Fig. 7e, Fig. S6**). This is likely the result of flagellar dysregulation by overactive PhoPQ (which inhibits the flagellar regulon in *Salmonella*; (Fàbrega and Vila, 2013)) in the absence of MgrB.

In the comparative transcriptomics data (**Fig. 7b**), we also noted a strong downregulation of the CsrB sRNA in Δ*mgrB* bacteria. In Enterobacteriaceae, CsrB titrates the global mRNA-binding protein and translational repressor CsrA through its 21 high-affinity CsrA binding sites (Vakulskas et al., 2015). We validated the downregulation of CsrB in Δ*mgrB* compared to wild-type *Salmonella* (log_2_FC = −3.37) by Northern blot (**Fig. 7c**). Rifampicin time course experiments suggested that lower CsrB levels in Δ*mgrB Salmonella* are likely the result of reduced sRNA stability in the absence of MgrB (**Fig. S7**). Since MgrB is unlikely to directly interact with cellular RNAs and CsrB has not been previously identified as a member of the PhoPQ regulon, further work will be necessary to dissect the effect of MgrB besides PhoQ.

### Proteins affected by MgrB deficiency include TCSs and effector proteins

Next, we integrated the comparative transcriptomics with proteomics data from the wild-type, Δ*mgrB,* and *mgrB*^+^ *Salmonella* strains grown under the SPI-2-inducing condition (**Table S3**; the overlap of the transcriptomic and proteomic datasets is shown in **Fig. 7f**). The levels of 58 proteins were significantly altered (*p*-value < 0.01 based on all possible permutations) in Δ*mgrB* relative to wild-type and *mgrB*^+^ *Salmonella*. To prioritize identification of MgrB-specific effects and given that *mgrB* levels are elevated in the complementation strain over wild-type *Salmonella* (**Fig. 7c**), we focused on those proteins that were dysregulated in both Δ*mgrB* and *mgrB*^+^ relative to wild-type, but with opposite expression patterns.

Of the resulting 58 proteins, 31 were accumulated in Δ*mgrB* and included the four T3SS effectors SlrP, SspH2, SrfA, and SopB (SopB dysregulation was validated in **Fig. 7d**) and members of four TCSs (SsrB, PhoB, RstA, and PhoP). Despite higher levels of PhoP protein, we did not observe increased *phoP* transcript levels in Δ*mgrB* in the above RNA-seq data (**Fig. 7b, Table S3**). Instead, higher PhoP protein levels in the *mgrB* mutant strain compared to the wild-type strain might be due to an increased translation rate of *phoP* mRNA or increased protein stability in the absence of this small protein. Conversely, 27 proteins were depleted in Δ*mgrB* compared to both wild-type and *mgrB*^+^ *Salmonella*. Among them were the motility proteins FliC, CheB, and GdhA, which were also repressed at the mRNA level (**Fig. 7b**).

Finally, in an attempt to disentangle the above molecular changes in Δ*mgrB Salmonella* from PhoP-dependent effects of MgrB, we consulted the SalCom Regulon database (Colgan et al., 2016). This resource contains RNA-seq data from several *Salmonella* mutants, each devoid of a single global transcriptional regulator or TCS, including a *phoP*-deficient mutant. Comparing the set of dysregulated genes in the transcriptomes of Δ*mgrB* and Δ*phoP Salmonella* revealed large overlaps, but in opposite directions. In the majority of cases, downregulation in Δ*mgrB* corresponded to an upregulation in Δ*phoP* and *vice versa* (examples are shown in **Fig. S8**). However, other TCSs were affected at the protein level in the absence of MgrB, but their expression was unaffected in Δ*phoP Salmonella*, indicating either an additional posttranscriptional regulation affecting their protein amount or a Δ*mgrB* effect which is independent of PhoPQ. Further efforts will be needed to identify alternative direct interaction partners of MgrB besides PhoQ.

## DISCUSSION

It is difficult to overstate the impact of post-genomic technologies on microbiology. It is now often as easy to measure a given functional parameter across the whole genome as it is to assay a single locus. The steady accumulation of these genome-wide datasets presents an opportunity for explorative analyses that integrate this information to produce testable hypotheses. We have taken this approach to small protein discovery and characterization in the model pathogen *Salmonella* Typhimurium, combining new purpose-generated datasets with those from previous studies to identify promising leads for further characterization.

### Refinement of *Salmonella* sORF annotation and prioritizing sORFs for functional characterization

To enrich the annotation of the *Salmonella* small proteome, we combined genome-based predictions by sPepFinder (Li and Chao, 2020) with ribosome profiling-based data from *Salmonella* grown in well-established conditions mimicking specific stages of the infection cycle (host cell invasion and intracellular replication). In this way, we identified 250 small proteins that are currently not present in the UniProt or *Salmonella* SL1344 (Kröger et al., 2013) annotations with experimental evidence for expression. In our predictions, 5’UTR-encoded sORFs, or “upstream open reading frames” (uORFs; 70 out of 250), and novel independently-transcribed sORFs (67 out of 250) were over-represented. The 5’UTR-encoded ORFs might be novel leader peptides or small proteins involved in the regulation of the proteins encoded downstream in the operon (Orr et al., 2019). Furthermore, out-of-frame sORFs, or alt-ORFs (30 out of 250), are an emerging subclass of small proteins with functional characterization lagging behind even “classical” sORFs (Meydan et al., 2019; Meydan et al., 2018; Weaver et al., 2019). Translation of three of the new sORFs, i.e. STsORF70, STsORF105, and STsORF44, the latter of which corresponds to *E. coli* MntS (Martin, 2011), was confirmed by Western blot. Future work will be necessary to validate translation of the remaining candidates, for example STsORF55, encoded in the 3’UTR of *pipC* (a.k.a *sigE*). Furthermore, as sPepFinder predicts sORFs with only AUG and GUG start codons, expanding the prediction to rare start codons might increase the number of candidate sORFs.

Another Ribo-seq study (Baek et al., 2017) undertook the effort of annotating novel small proteins in *Salmonella*, albeit in a different strain (*Salmonella* Typhimurium 14028s). Among the 80 sORF candidates that are conserved between the two *Salmonella* strains and have a size between 10 and 99 aa, only 16 were called by our approach (**Table S1**). This limited overlap might be caused by a combination of experimental and analytical differences between the two studies, or strain-specific expression differences, and future work will be required to assess the false-positive and false-negative rates in either approach.

In order to prioritize sORF candidates for functional characterization, we analysed both existing and newly generated global datasets with a specific focus on small proteins. For this, we merged our new sORFs with the previously annotated small proteome of *Salmonella*. 91 uncharacterized small proteins were detected in a Grad-seq dataset from *Salmonella* grown under SPI-1-inducing conditions. As some of these proteins are annotated solely based on homology to other bacterial species, their detection in our gradient represents the first direct evidence for their existence in *Salmonella*. Most of the detected small proteins migrated to low density fractions of the gradient, suggesting tight engagement in only small, if any, protein complexes. However, without specifically enriching for small proteins, upon digestion prior to mass spectrometry analysis, the relatively few peptides they give rise to are easily overwhelmed by the signal derived from average-sized proteins. Therefore, future modifications of the mass spectrometry approach used for Grad-seq, e.g. bypassing the fragmentation step, might further increase the detection of small proteins.

To identify virulence-related small protein candidates in *Salmonella*, we searched dual RNA-seq data (Westermann et al., 2016) for sORFs induced during the infection of epithelial cells. A highly induced sORF during infection was *yjiS*, encoding an uncharacterized small protein with a DUF1127 domain. Likewise, analysis of TraDIS fitness data from *Salmonella* transposon mutants in a macrophage infection model suggested YjiS as a new anti-virulence factor in *Salmonella*. These findings propose YjiS as a high-priority sORF candidate for future characterization.

### The small protein MgrB contributes to *Salmonella* virulence

MgrB was one of the few small proteins with a previously investigated molecular function in *Salmonella*. MgrB regulates the activity of the PhoPQ TCS (Lippa and Goulian, 2009), a function conserved in *E. coli* (Salazar et al., 2016). Here, we showed for the first time that, relative to the isogenic wild-type strain, Δ*mgrB Salmonella* is impaired at all stages of macrophage and epithelial cell infection. We uncovered a positive effect of MgrB on flagella and motility-related genes at both the transcript and protein level. Upon further inspection of the data, we could link this effect not only to PhoPQ, long known to repress flagella expression (Adams et al., 2001), but also to the TCS SsrAB, the master regulator of the SPI-2 regulon (Walthers et al., 2007). Among others, the transcription factor SsrA is responsible for repressing *flhDC*, encoding the master regulators of the flagellar expression cascade (Ilyas et al., 2018). In our proteomics data, the histidine kinase SsrB was upregulated in the Δ*mgrB* mutant. This could contribute to the observed inhibition of flagellar gene expression in the absence of MgrB. Furthermore, elevated SsrB levels are known to lead to a defect in epithelial cell invasion (Pérez-Morales et al., 2017), another phenotype we found associated with the lack of MgrB.

MgrB also affected the expression of numerous genes that are currently not considered members of the PhoPQ regulon. This suggests additional, PhoPQ-independent roles for MgrB, supported by recent findings of *E. coli* MgrB interacting with further histidine kinases such as PhoR (Yadavalli et al., 2018). PhoB, the response regulator of the PhoBR TCS, was also detected among the upregulated proteins in our Δ*mgrB* mutant. As histidine kinases and response regulators of different TCSs share several conserved domains, crosstalk between such systems has been hypothesized (Laub and Goulian, 2007). It is therefore tempting to speculate that MgrB could act as a regulator of different histidine kinases. We note, however, that the dissection of PhoP-dependent and PhoP-independent MgrB effects is hampered by the fact that *mgrB* is itself a member of the PhoPQ regulon. Uncoupling PhoPQ-dependent from PhoPQ-independent effects will be necessary to further asses the role(s) of MgrB in *Salmonella* virulence.

## CONCLUSIONS

Our integrative approach to identifying and prioritizing small proteins for further study in *Salmonella* will serve as a blueprint for other species. Countless global datasets are available for diverse bacterial organisms including Gram-positive species. For example, high-resolution transcriptomics, transposon mutagenesis, and Grad-seq data exist for *Streptococcus pneumoniae* (Aprianto et al., 2016, 2018; Hor et al., 2020; Rowe et al., 2019; van Opijnen and Camilli, 2012; Warrier et al., 2018). More generally, various transposon-sequencing approaches have been applied to bacterial pathogens under virulence conditions (Karlinsey et al., 2019; Rendón et al., 2020; Warr et al., 2019), and dual RNA-seq has become the gold-standard to chart the transcriptional landscape of pathogens during infection (Montoya et al., 2019; Pisu et al., 2020; Ritchie and Evans, 2019; Westermann et al., 2017). Re-inspection of these data may provide invaluable information on potentially new biological roles carried out by small proteins in bacterial pathogenesis.

---

**Supplementary Figure S1:**
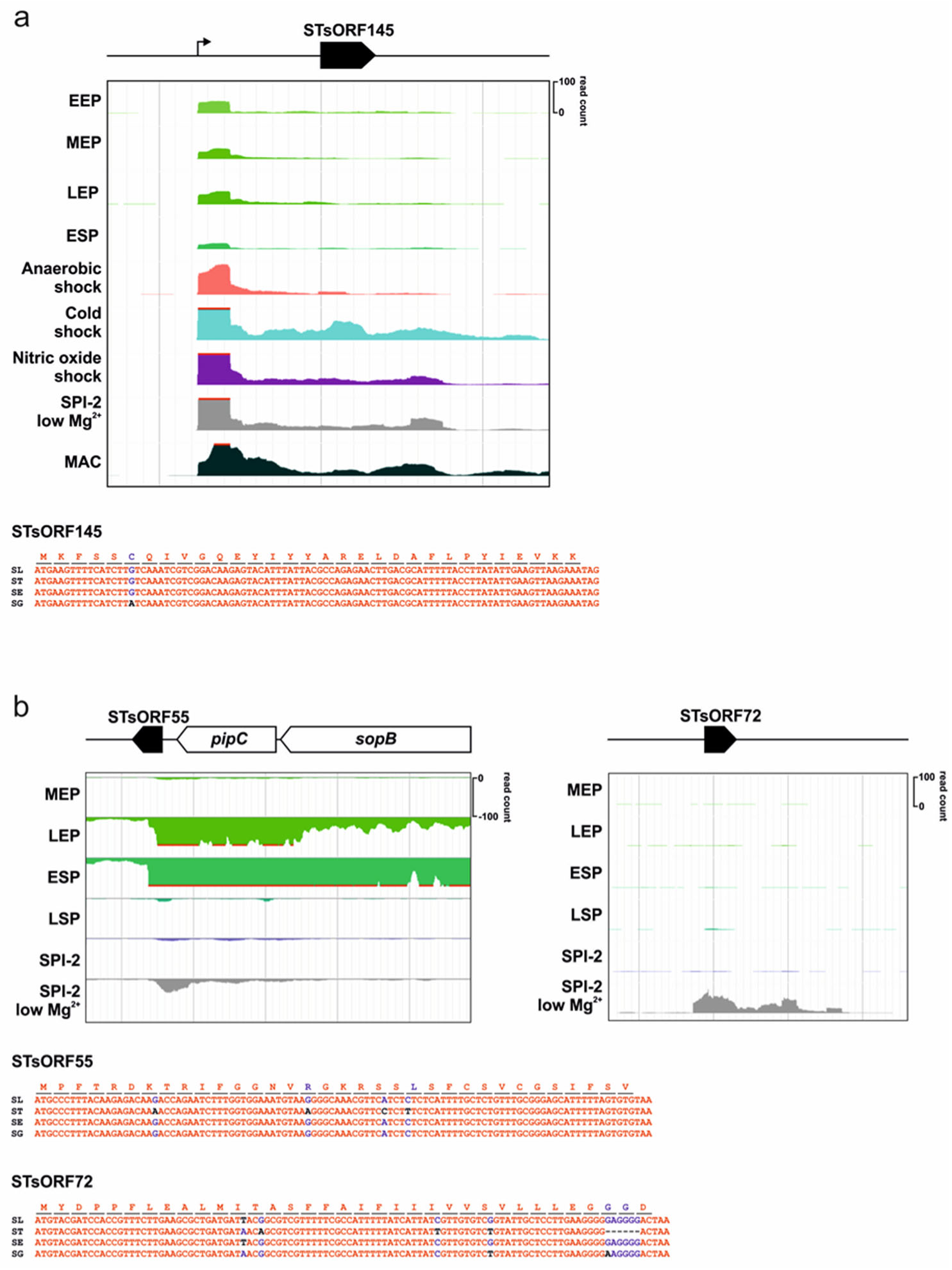
Annotation and expression of STsORFs with condition-specific expression patterns. **a**, STsORF145 is upregulated in response to different shocks. **b**, STsORF55 (left), encoded in the 3’UTR of *pipC*, is expressed in SPI-1-inducing conditions while the STsORF72 (right), for which no TSS has been annotated, in expressed in SPI-2-inducing conditions with low Mg^2+^. In both panels the genomic location of the new sORFs (black-filled arrows) relative to annotated genes and annotated TSSs (from (Kröger et al., 2013), thin arrows) are shown, as well as their expression patterns from SalComMac (Srikumar et al., 2015), and their conservation among *Salmonellae*. EEP: early exponential phase; MEP: mid-exponential phase; LEP: late exponential phase; ESP: early stationary phase; LSP: late stationary phase; SPI-2: SPI-2-inducing; MAC: inside macrophages. Alignments of the CDSs from the indicated *Salmonellae* (*Salmonella* Typhimurium: “SL”; *Salmonella* Typhi: “ST”; *Salmonella enteritidis*: “SE”; *Salmonella* Gallinarum: “SG”). On top, the corresponding amino acid sequence is shown. *Salmonella bongori* was excluded from all the alignments as lacking conservation at the nucleotide level. Nucleotides and amino acids are coloured in red if the conservation is 100%, in blue if it is higher than 90%, and in black if it is less than 10%.

**Supplementary Figure S2:**
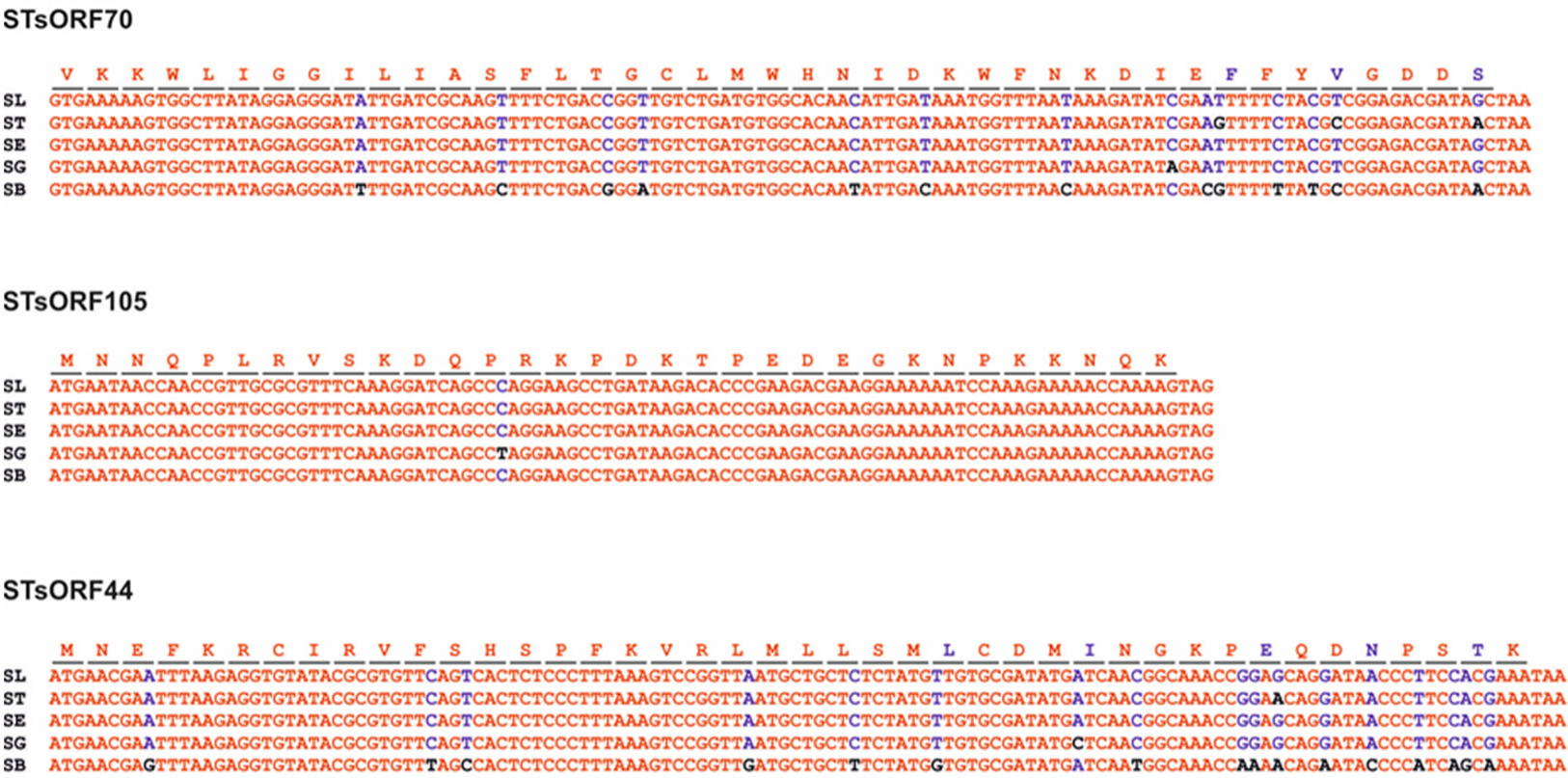
Validated STsORFs conservation. The alignments show the conservation of the three validated sORFs among different *Salmonellae*. *Salmonella* Typhimurium: “SL”; *Salmonella* Typhi: “ST”; *Salmonella enteritidis*: “SE”; *Salmonella* Gallinarum: “SG”; *Salmonella bongori*: “SB”. Nucleotides are coloured in red if the conservation is 100%, in blue if it is higher than 90%, and in black if it is less than 10%. On top is reported the corresponding amino acid sequence.

**Supplementary Figure S3:**
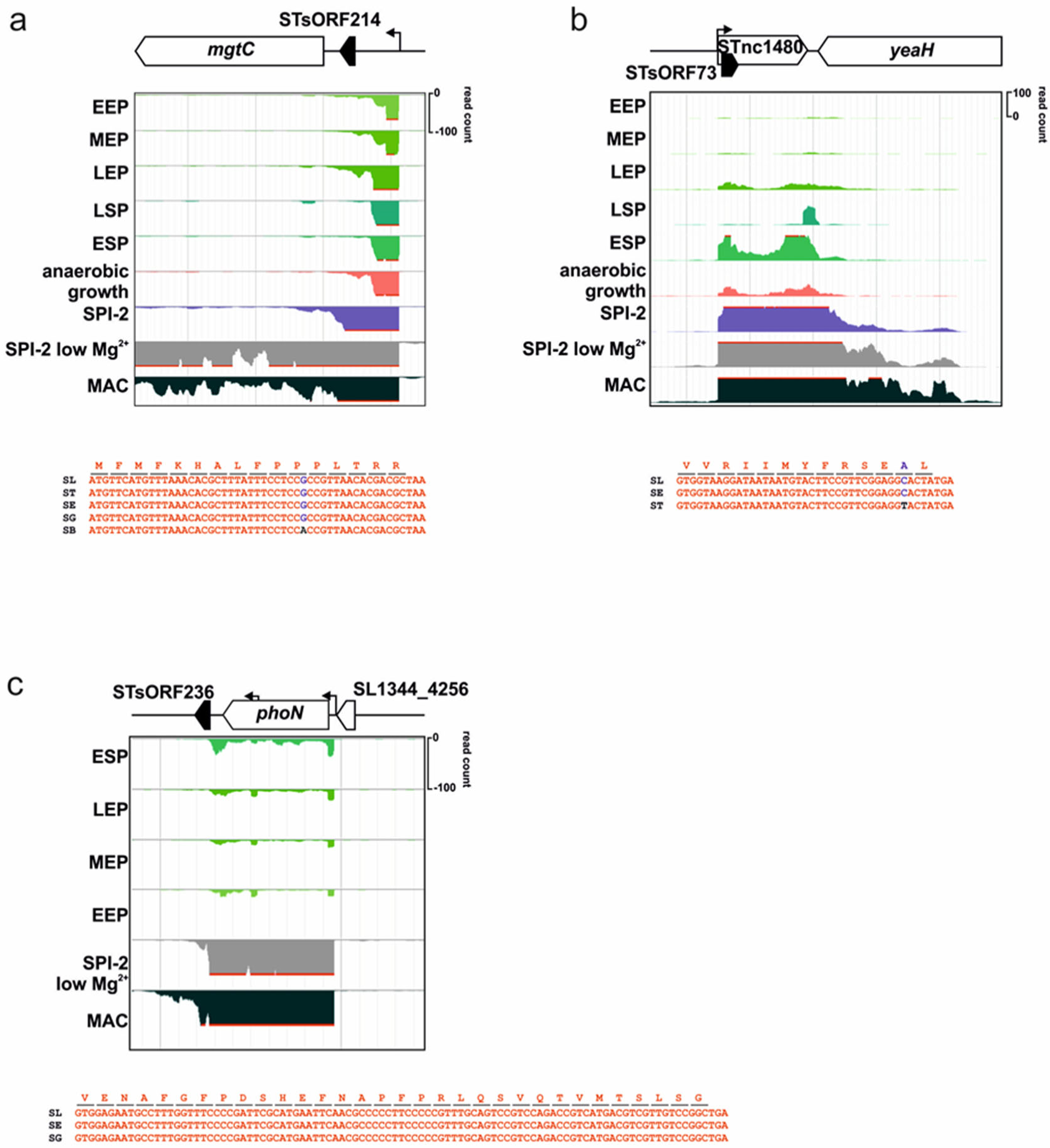
Genome localization and expression pattern of STsORFs highly induced in infection. SalComMac (Srikumar et al., 2015) expression patterns of **a**, STsORF214, **b**, STsORF73, and **c**, STsORF236 relative to annotated genes and TSSs (from (Kröger et al., 2013)). Growth conditions: EEP: early exponential phase; MEP: mid exponential phase; LEP: late exponential phase; LSP: late stationary phase; ESP: early stationary phase; MAC: macrophages. Below are reported the amino acid sequence and the alignments of the two CDSs among *Salmonella* Typhimurium: “SL”; *Salmonella* Typhi: “ST”; *Salmonella enteritidis*: “SE”; *Salmonella* Gallinarum: “SG”. The strains reported are those where the sequence was conserved.

**Supplementary Figure S4:**
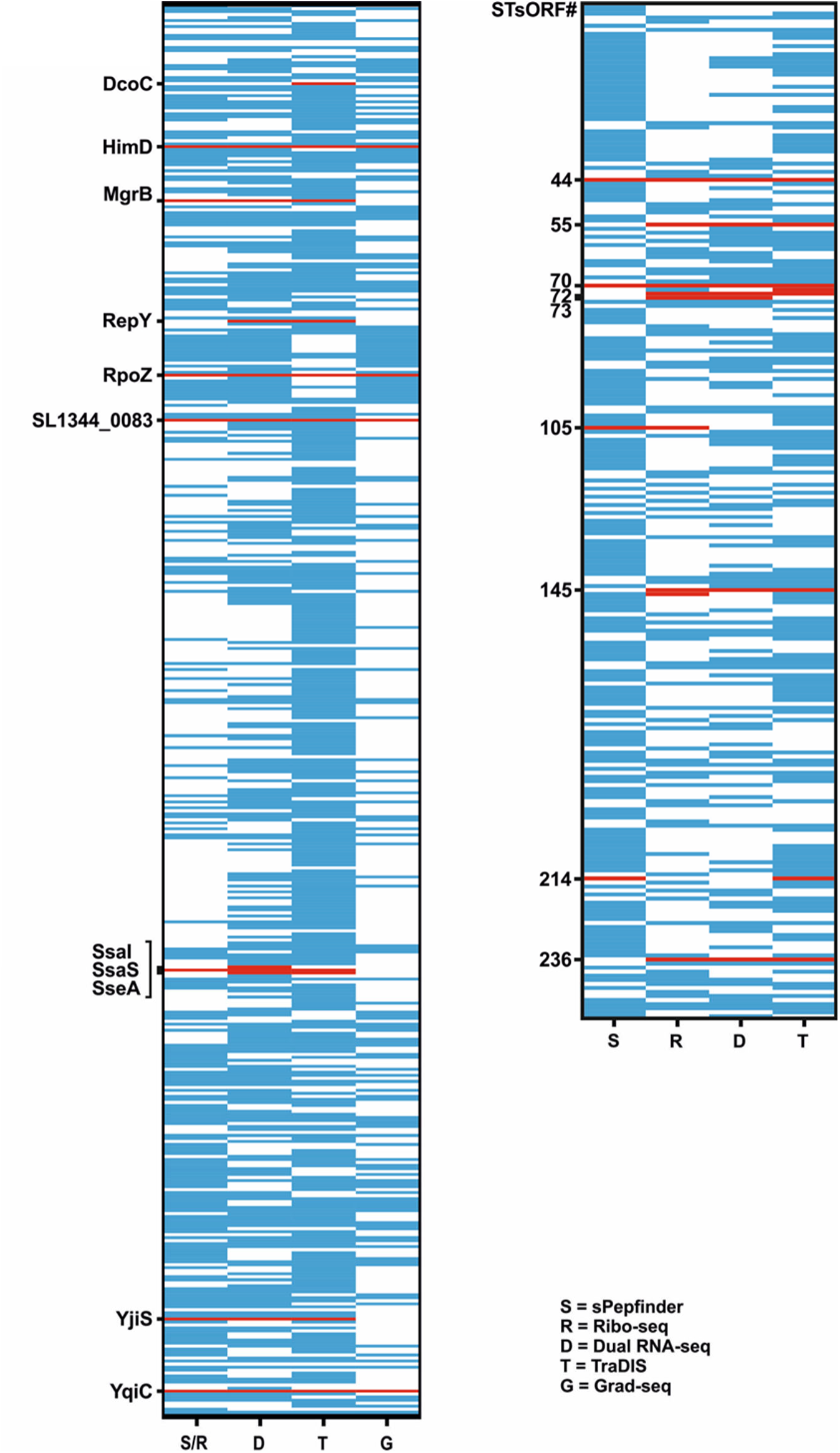
Overview of small proteins occurrence in the analysed datasets. Global map of the newly predicted small proteins (250, shorter than 100 aa) and the currently annotated sORFs (< 100 aa) indicating the dataset in which they were detected (blue box). “S” = sPepFinder and Ribo-seq; “D” = dual RNA-seq; “T” = TraDIS; “G” = Grad-seq. For Ribo-seq, sORFs are considered detected if their translational efficiency (70S footprint/total RNA) is > 0.5 in at least one condition. For TraDIS, all sORFs targeted by transposon insertion were considered. The bottom panel is the enlarged map of the 250 novel small proteins identified in this study. The red boxes (substituting blue boxes) highlight proteins of interest mentioned throughout the text.

**Supplementary Figure S5:**
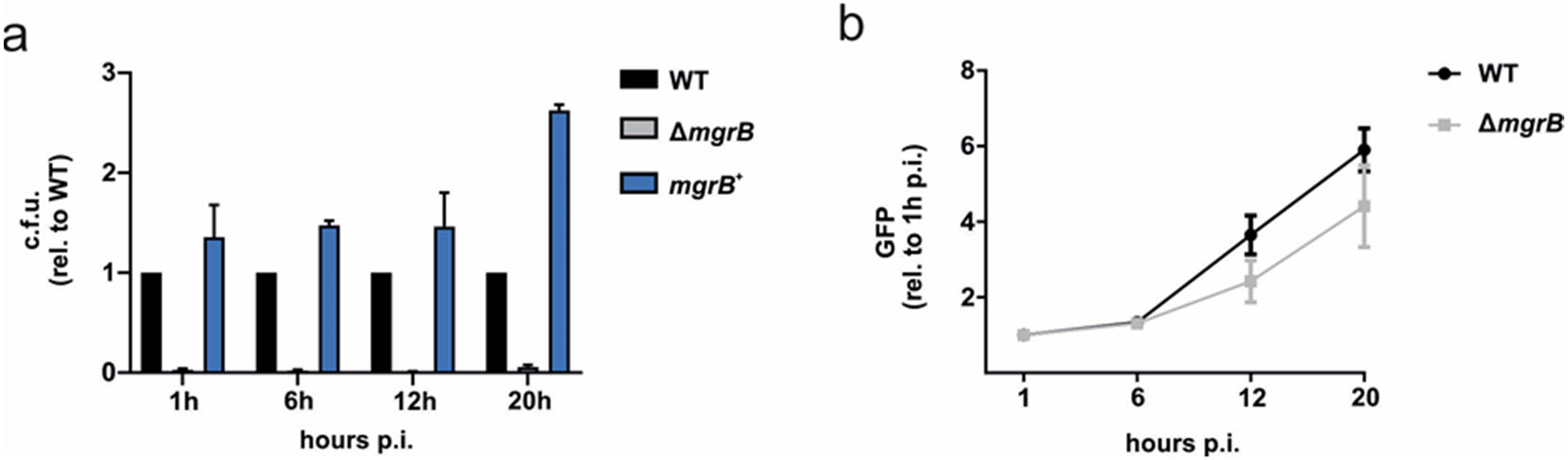
The requirement of MgrB for epithelial cell infection. **a**, Gentamicin protection assay of *Salmonella* wild-type, Δ*mgrB* or *mgrB^+^* infecting HeLa cells. Data were collected from three biological replicates with error bars indicating standard deviations from the mean. **b**, Intracellular replication in HeLa cells of *Salmonella* wild-type or Δ*mgrB* expressing GFP. Fluorescence was normalized to the 1 h time point. Data were collected from three biological replicates with error bars indicating standard deviations from the mean.

**Supplementary Figure S6:**
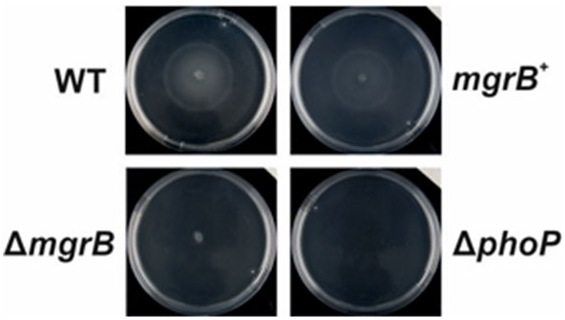
Impact of MgrB on *Salmonella* motility. The swimming defect of *Salmonella* Δ*mgrB* compared to the wild-type strain is restored in the complementation strain *mgrB^+^*. The Δ*phoP* strain was included as a negative control. Images are representative of three biological replicates.

**Supplementary Figure S7:**
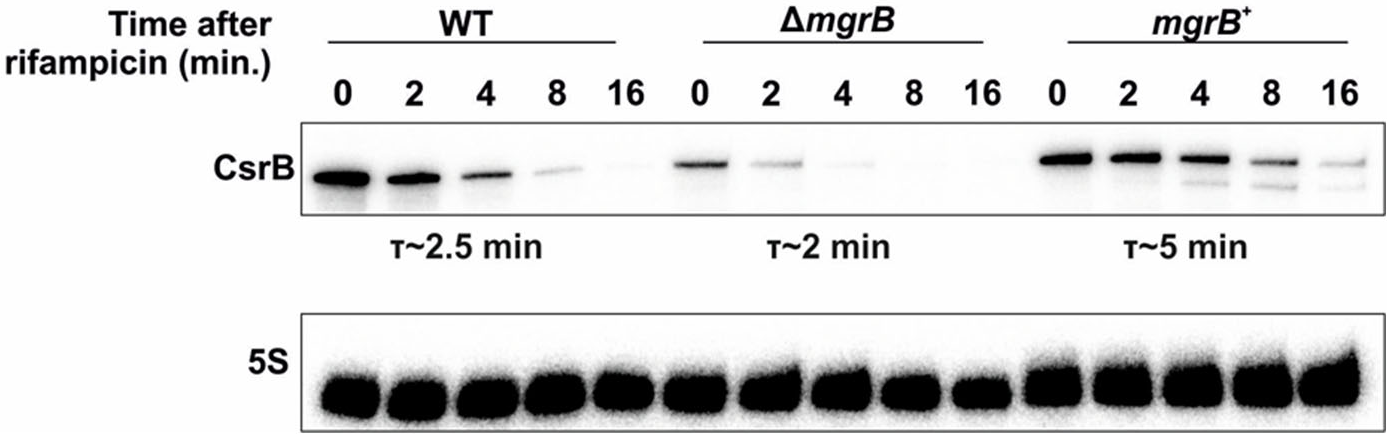
CsrB half-life is shorter in the absence of MgrB. Rifampicin assay of CsrB stability in *Salmonella* wild-type, Δ*mgrB*, and *mgrB^+^*. Samples for total RNA were collected at 0, 2, 4, 8, and 16 minutes after rifampicin addition. CsrB levels were analysed via Northern Blot. Quantification of the half-lives was calculated averaging three biological replicates.

**Supplementary Figure S8:**
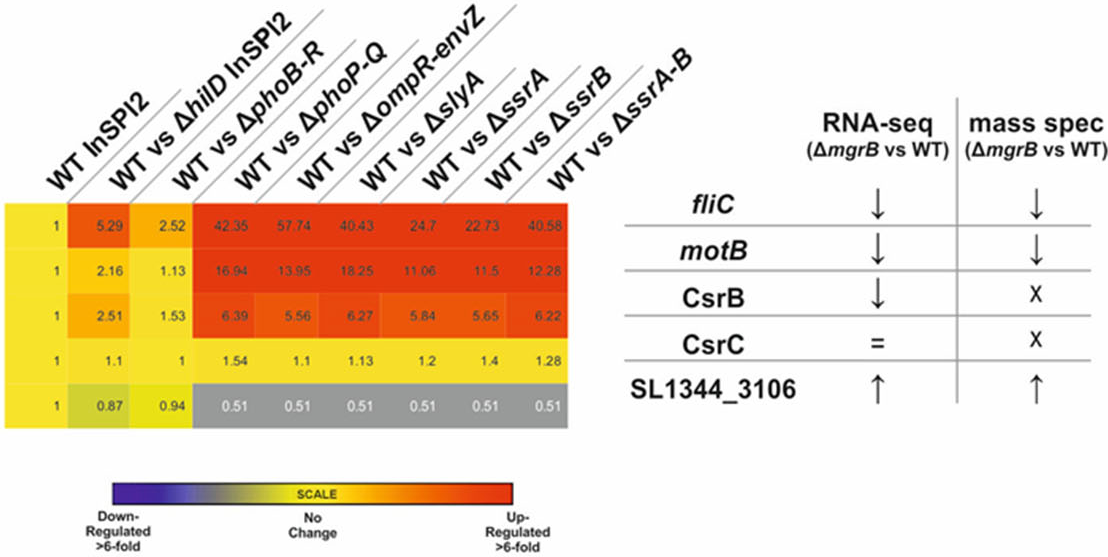
Expression patterns of MgrB-regulated transcripts in different *Salmonella* deletion mutants. Transcriptomic data of, among others, Δ*phoPQ Salmonella* grown under SPI-2-inducing conditions, taken from SalComRegulon (Colgan et al., 2016). The numbers indicate fold-change in abundance relative to the wild-type strain grown in the same condition. These data show the upregulation of transcripts that are downregulated in our datasets (and vice versa) in *Salmonella* Δ*mgrB* vs the wild-type strain.

## LIST OF SUPPLEMENTARY TABLES

**Supplementary Table S1:** Excel file with sPepFinder-predicted and Ribo-seq-predicted sORFs.

**Supplementary Table S2** Excel file with data from the re-analysis of dual RNA-seq time course (Westermann et al., 2016), including predicted sORFs from sPepFinder and Ribo-seq. Data from TraDIS infection. Sedimentation profile of small proteins detected in Grad-seq (Gerovac *et al*., submitted).

**Supplementary Table S3:** Excel file with RNA-seq data of *Salmonella* wild-type vs Δ*mgrB* grown *in vitro* in SPI-2 medium and mass spectrometry data of *Salmonella* wild-type, Δ*mgrB*, and *mgrB*^+^ grown *in vitro* in SPI-2 medium.

**Supplementary Table S4:** Excel file with list of strains, oligonucleotides, and antibodies used in this work.

## METHODS

### Strains and growth conditions

All bacterial strains and plasmids used for this study are reported in the **Table S4**. The Δ*mgrB* strain, as well as the SPA-tagged strains, were generated as previously described (Datsenko and Wanner, 2000). Oligonucleotides used for cloning can be found in **Table S4**. The respective mutations were then transduced in the wild-type or GFP^+^ background using P22 phages (Sternberg and Maurer, 1991). For routine growth of *Salmonella*, a single bacterial colony was grown overnight in Lennox-Broth (LB) medium at 37°C with shaking at 220 rpm, diluted 1:100 into fresh medium, and then grown to the required cell density. The SPI-1-inducing condition is defined as growth in LB to an OD_600_ of 2.0 (Kröger et al., 2013). For growth in SPI-2-inducing conditions (Löber et al., 2006), cells that had reached SPI-1 conditions were centrifuged for 2 min at 12,000 rpm at room temperature (RT), washed twice with PBS (Gibco) and once with SPI-2 medium (Löber et al., 2006), and then diluted 1:50 into fresh SPI-2 medium. The cultures were again grown at 37°C and 220 rpm to an OD_600_ 0.3. When required, the medium was supplemented with 100 μg/ml ampicillin.

### Mammalian cell cultures

HeLa-S3 cells (ATCC CCL-2.2) and RAW.B mouse macrophages (RAW264.7; ATCC TIB-71) were cultured as described in (Westermann et al., 2016). HeLa cells were passaged in DMEM (Gibco) and RAW.B cells were passaged in RPMI (Gibco) medium supplemented with 10% fetal calf serum (FCS, Biochrom), 2 mM L-glutamine (Gibco) and 1 mM sodium pyruvate (Gibco) in T-75 flasks (Corning). Cells were grown in a 5% CO_2_, humidified atmosphere at 37°C, and routinely tested for mycoplasma contamination with the MycoAlert Mycoplasma Detection kit (Lonza). Two days before infection, 2 x 10^5^ cells were seeded in six-wells plates (2 ml medium).

### sPepFinder

sPepFinder is a support vector machine learning-based computational approach for *ab initio* prediction of bacterial small ORFs (Li and Chao, 2020). Briefly, it first extracts informative features from a collection of sequence and structural properties of known bacterial small proteins. The informative features include a thermodynamics model of ribosome binding sites and amino acid composition. sPepFinder has achieved a 92.8% accuracy in 10-fold cross validation in a test dataset of ten bacterial species (eight from the Enterobacteriaceae family, as well as *Vibrio cholerae* and *Pseudomonas*).

### Ribosome profiling

Ribosome profiling was performed as previously described (Oh et al., 2011) with modifications. *Salmonella* wild-type was grown in LB, SPI-1- and SPI-2-inducing conditions as described above. Bacterial cells were harvested from cultures by fast-filtration with a 0.45 µm polyethersulfone membrane (Millipore) and immediately frozen in liquid N_2_. Before harvesting, a sample was taken for total RNA analysis, mixed with 0.2 vol stop mix (5% buffer-saturated phenol (Roth) in 95% ethanol). Cell pellets were resuspended in ice-cold lysis buffer (100 mM NH_4_Cl, 10 mM MgCl_2_, 20 mM Tris-HCl, pH 8.0, 0.1% NP-40, 0.4% Triton X-100 (Gibco), 50 U/ml DNase I (Fermentas), 500 U RNase Inhibitor (moloX, Berlin), 1 mM chloramphenicol) and lysed using glass beads and vortexing at high speed for 10 x 30 s, with chilling on ice in between each round. Lysates were clarified by centrifugation at 21,000*g* for 10 min. Next, 14-17 A_260_ of lysate was digested with 800 U/A_260_ of micrococcal nuclease (MNase, NEB) at 25°C with shaking at 14,500 rpm for 20 min in lysis buffer supplemented with 2 mM CaCl_2_ and 500 U RNase Inhibitor. A mock-digested control (no enzyme added) was also included for each lysate to confirm the presence of polysomes. Digests were stopped with ethylene glycol-bis(β-aminoethyl ether)-*N*,*N*,*N*′,*N*′-tetraacetic acid (EGTA, final concentration 6 mM) and immediately loaded onto 10 – 55% sucrose gradients prepared in sucrose buffer (100 mM NH_4_Cl, 10 mM MgCl_2_, 5 mM CaCl_2_, 20 mM Tris-HCl, pH 8.0, 1 mM chloramphenicol) with 2 mM fresh dithiothreitol. Gradients were centrifuged in a SW40-Ti rotor in a Beckman Coulter Optima L-80 XP ultracentrifuge for 2h 30 min at 35,000 rpm at 4°C, and then 70S monosome fractions were collected using a Gradient Station *ip* (Biocomp Instruments). RNA was extracted from fractions or cell pellets for total RNA using hot phenol-chloroform-isoamyl alcohol (25:24:1, Roth) or hot phenol, respectively, as described previously (Sharma et al., 2007; Vasquez et al., 2014). Ribosomal RNA was depleted from 5 µg of total RNA by subtractive hybridization with a complex probe set for *Salmonella enterica* (Senterica_riboPOOL-RP1, siTOOLs, Germany) according to the manufacturer’s protocol with Dynabeads MyOne Streptavidin T1 beads (Invitrogen). Total RNA was fragmented with RNA Fragmentation Reagent (Ambion). Monosome RNA and fragmented total RNA was size-selected (26-34 nt) on 15% polyacrylamide/7 M urea gels as described previously (Ingolia et al., 2012) using RNA oligonucleotides NI-19 and NI-20 as guides. RNA was cleaned up and concentrated by isopropanol precipitation with 15 μg GlycoBlue (Ambion) and dissolved in H_2_O. Libraries were prepared by Vertis Biotechnologie AG (Freising, Germany) using a small RNA protocol without fragmentation and sequenced on a NextSeq500 instrument (high-output, 75 cycles) at the Core Unit SysMed at the University of Würzburg.

### Analysis of ribosome profiling data

Read files were processed and analyzed with HRIBO (version 1.4.3)(Gelhausen et al., 2020), a snakemake (Köster and Rahmann, 2012) based workflow that downloads all required tools from bioconda (Grüning et al., 2018) and automatically determines the necessary processing steps. We additionally use pinned tool versions which ensures reproducibility of the analysis. Adapters were trimmed from the reads with cutadapt (version 2.1) (Martin, 2011) and then mapped with segemehl (version 0.3.4) (Otto et al., 2014). Reads mapping to ribosomal RNA genes and multi-mappers were removed with SAMtools (version 1.9) (Li et al., 2009) using the rRNA annotation. Open reading frames were called with an adapted variant of REPARATION (Ndah et al., 2017) which uses blast instead of usearch (see https://github.com/RickGelhausen/REPARATION_blast). Quality control was performed by creating read count statistics for each processing step and RNA-class with Subread featureCounts(1.6.3) (Liao et al., 2014). All processing steps were analyzed with FastQC (version 0.11.8) (Andrews) and results were aggregated with MultiQC (version 1.7) (Ewels et al., 2016). Summary statistics for all available annotated and merged novel ORFs detected by REPARATION were computed in a tabularized form including translational efficiency, RPKM normalized readcounts, codon counts, nucleotide and amino acid sequences, etc. Additionally, GFF track files with the same information were created for in-depth genome browser inspection, in addition to GFF files showing potential start/stop codon and RBS information.

### Dual RNA-seq

Data were taken from (Westermann et al., 2016). In brief, GFP+ *Salmonella* was used to infect HeLa cells (HeLa-S3; ATCC CCL-2.2) with an MOI (multiplicity of infection) of 5. At different time points (2, 4, 8, 16, 24 h p.i.) cells were collected and sorted to enrich for infected epithelial cells (GFP+). These cells were subjected to RNA extraction and sequencing after rRNA depletion. RNA sequencing was also performed on *Salmonella* grown in LB to OD_600_ 2.0, which represents the inoculum used for infection. Re-analysis of the data was carried out as in (Westermann et al., 2016) with our updated annotation.

### TraDIS

A *Salmonella* transposon mutant library containing circa 100,000 mutants was generated using EZ-Tn5 transposase (Epicentre) and the *aphA1* kanamycin resistance gene as described previously (Langridge et al., 2009). Two days before infection, RAW.B cells were seeded at a density of 2 x 10^6^ in two 75 ml flasks per replicate in RPMI supplemented with penicillin and streptomycin, and then changed to an antibiotic free medium one day before infection. An aliquot of 1 OD_600_/ml equivalent of the *Salmonella* library was grown in 200 ml LB with 10 μg/ml kanamycin at 37°C overnight with shaking. Next, 2 OD_600_ equivalents of this overnight culture were pelleted and resuspended in RPMI with 10% mouse serum for 20 min at room temperature (RT) for opsonization. A similar amount of overnight culture was pelleted and used for input genomic DNA preparation. The RAW.B cells were then infected directly in flasks at an MOI of 20, centrifuged for 10 min at 250*g* at RT and incubated for 30 min at 37°C. The medium was then replaced with RPMI containing 100 μg/ml gentamicin, incubated for an additional for 30 minutes, then washed with PBS and replaced again with RPMI containing 10 μg/ml gentamicin for the remaining duration of the experiment. At 20 h p.i., the medium was aspirated, and cells were washed once with PBS before being scraped from the flask in 10 ml PBS and harvested by centrifugation for 10 min at 250 *g.* The supernatant was discarded, and samples for each replicate were pooled in 6 ml PBS containing 0.1% (v/v) Triton X-100. This was incubated for 10 min at RT with occasional vortexing, before being centrifuged again at 250*g* for 10 min. The supernatant was recovered, pelleted, and used for DNA extraction. DNA was extracted using the phenol-chloroform method. Briefly, bacterial pellets were resuspended in 250 μl of a 50 mM Tris-HCl, 50 mM EDTA, pH 8 solution and frozen for at least 1 hour at −20°C. Pellets were then defrosted at RT and treated with 2.5 μg/ml lysozyme on ice for 45 min, followed by 2.4 μg/ml per OD unit of input culture RNase A (Fermentas) for 40 min at 37°C, and then ∼333 μg/ml proteinase K in buffer (0.5% (w/v) SDS, 50 mM Tris-HCl, 0.4 M EDTA, pH 8) at 50°C until the sample cleared (approx. 30 min – 1h). This solution was then mixed with 300 μl milliQ filtered water and added to a phase lock gel (PLG) tube containing 400 μl phenol/chloroform (Roth). The sample was vigorously mixed by inversion, then centrifuged at 15°C for 15 minutes. The aqueous phase was collected, then precipitated with 1.4 ml 100% ethanol containing 0.1 M sodium acetate and inverted 6-8 times. This solution was then centrifuged at 13,000 rpm at RT for 20 min, the supernatant was discarded, and the pellet was washed with 70% ethanol before drying and resuspension in milliQ filtered water. Sequencing of 2 replicate infection experiments was performed at the Wellcome Trust Sanger Institute using the TraDIS dark-cycle sequencing protocol for 50 cycles on a MiSeq sequencer (Illumina) with a read count yield of between 1.3 and 1.9 million reads as previously described (Barquist et al., 2016). The reads were then processed with the Bio-TraDIS Toolkit (https://github.com/sanger-pathogens/Bio-Tradis, (Barquist et al., 2016)), with ∼98-99% of reads matching the expected transposon tag sequence, and subsequent read mapping rates of ∼94-98%. Transposon read counts per gene were then summarized using the tradis_gene_insert_sites script, excluding insertions in the first and last 10% of each gene. Infected samples were then compared to controls using the tradis_comparison.R script, using edgeR (Robinson et al., 2010), and filtered for genes with a |log_2_FC| > 1 and a q-value < 0.05.

### Grad-seq

Gradient profiling mass spectrometry data were taken from (Gerovac *et al*., submitted), based on the Grad-seq approach first described in (Smirnov et al., 2016). In brief, *Salmonella* wild-type was grown in LB to an OD_600_ of 2.0, collected, and lysed by glass bead beating. The lysate was then loaded on a linear glycerol gradient (10-40% w/v) and separated by ultracentrifugation. The gradient was then fractionated and each fraction, including the pellet, was analysed by mass spectrometry.

### RNA extraction

For total RNA preparation, cells were grown in SPI-2 medium to an OD_600_ of 0.3 as previously described. Then, 10 ml of culture were mixed with 2 ml of STOP solution (95% ethanol, 5% phenol), snap frozen in liquid nitrogen and total RNA was extracted via the “hot-phenol” method (Vasquez et al., 2014). Briefly, the frozen culture was thawed, centrifuged 20 minutes at 4,500 rpm and 4°C and the cell pellet resuspended in 0.5 mg/ml lysozyme in TE, pH 8.0. Next, 60 μl of 10% w/v SDS was added, the tube was mixed by inversion, and incubated for 2 min at 64°C. After this, 66 μl of 3 M sodium acetate, pH 5.2 was added, followed by 750 μl phenol (Roti Aqua phenol, Roth). The solutions were mixed by inversion and incubated 6 min at 64°C. Tubes were then cooled on ice and centrifuged 15 min at 13,000 rpm at 4°C. The aqueous phase was then moved to 2 ml PLG tubes, to which 750 μl chloroform were added, shaken and centrifuged 12 min 13,000 rpm at 4°C. The aqueous phase was moved to new tubes to which were added 2 volumes of 30:1 mix (ethanol:3 M sodium acetate, pH 6.5). This was left to precipitate over night at −20°C, then centrifuged for 30 min at 4°C and 13,000 rpm, washed with 75% v/v ethanol, and resuspended in RNase-free water by shaking for 5 min at 65°C. To remove DNA contamination, samples were treated with 0.25 U of DNase I (Fermentas) for 1 μg RNA for 45 min at 37°C.

### Total RNA-seq and analysis

cDNA libraries of total RNA were prepared at Vertis Biotechnologie AG after rRNA depletion (Ribo-Zero rRNA Removal Kit (Bacteria); Illumina). Sequencing was performed on an Illumina NextSeq 500 platform with approximately 20 million reads per library. The adapters were removed from the FASTQ format reads using cutadapt, then quality trimming was carried out with fastq_quality_trimmer from the FastX suite (Version 0.3.7). Alignment to the *Salmonella* Typhimurium SL1344 genome downloaded from NCBI (NC_016810, NC_017718, NC_017719, and NC_017720) was performed using READemption (Förstner et al., 2014) (Version 0.4.5). Differential gene expression analysis was carried out with edgeR (Robinson et al., 2010) (Version 3.20.8). Only genes with at least 5 uniquely mapped reads in two experiments were considered. The cut-off for differential expression was set to q-value < 0.05 and |log_2_FC| > 2.

### Northern blot analysis

Northern blot analysis of 10 μg of RNA per sample was performed with 6% (v/v) polyacrylamide-7 M urea gels as previously described (Westermann et al., 2016). ^32^P-labelled DNA oligonucleotides (**Table S4**) complementary to the transcript of interest was then incubated in Hybri-Quick buffer (Carl Roth AG) on the membranes (Hybond XL membranes, Amersham) at 42°C overnight, followed by sequential washes in SSC buffers (5x, 1x, 0.5x) with 0.1% SDS. The membranes were then exposed on phosphor screen and revealed on a Typhoon FLA 7000 phosphorimager (GE Healthcare). Signal quantification was performed using ImageJ (Schneider et al., 2012).

### Western blotting

For Western blot validation of novel STsORFs and mass spectrometry data, cells were grown to the appropriate OD_600_ in SPI-1 or SPI-2 medium, harvested by centrifugation at RT for 2 min at 16,000*g,* and resuspended in 1x protein loading buffer (5x solution: 5g SDS, 46.95 ml Tris-HCl, pH 6.8, 0.075 g bromophenol blue, 75 ml glycerol, 11.56 g dithiothreitol, filled up to 150 ml H_2_O) at a concentration of 0.01 OD/μl. Samples were denatured for 5 min at 95°C and 0.1 OD equivalents were separated on 15% SDS-PAGE gels. Separated proteins were then transferred to a PVDF membrane (Perkin Elmer) for 90 min with a semidry blotter (Peqlab; 0.2 mA/cm^2^ at 4°C) in transfer buffer (25 mM Tris base, 190 mM glycine, 20% (v/v) methanol). After transfer, membranes were blocked for 1 h at RT in 10% (w/v) milk/TBS-T20 (Tris-buffered-saline-Tween-20). The membranes were then incubated with the appropriate primary antibody (**Table S4**) at 4°C overnight, and then after three 5 min washes with TBS-T20, incubated with the corresponding secondary antibody (**Table S4**) for 1 h at RT. At the end of the hybridization, the membranes were again washed three times for 5 min with TBS-T20 and the blots developed with Western lightning solution (Perkin Elmer) with a Fuji LAS-4000 imager (GE Healthcare).

### Whole proteome preparation

For the preparation of the total *Salmonella* proteome for mass spectrometry analysis, cells were grown in SPI-2-inducing conditions (see above). At an OD_600_ of 0.3, cells were pelleted, washed, and resuspended in protein loading dye for loading on a precast gel at a concentration of 1 OD/100 μl. Proteins were separated by 1D SDS-PAGE and prepared for MS/MS analyses as previously described (Bonn et al., 2014). Briefly, gel lanes were fractionated into 10 gel pieces, cut into smaller blocks and transferred into low-binding tubes. Samples were and washed until gel blocks were destained. After drying of gel pieces in a vacuum centrifuge, they were covered with trypsin solution. Digestion took place at 37 °C overnight before peptides were eluted in water by ultrasonication. The peptide-containing supernatant was transferred into a fresh tube, desiccated in a vacuum centrifuge and peptides were resolubilized in 0.1% (v/v) acetic acid for mass spectrometry analysis.

### MS/MS analysis

Tryptic peptides were subjected to liquid chromatography (LC) separation and electrospray ionization-based mass spectrometry applying exactly the same injected volumes in order to allow for label-free relative protein quantification. Therefore, peptides were loaded on a self-packed analytical column (OD 360 μm, ID 100 μm, length 20 cm) filled with of Reprosil-Gold 300 C18, 5 µm material (Dr. Maisch, Ammerbuch-Entringen, Germany) and eluted by a binary nonlinear gradient of 5 - 99% acetonitrile in 0.1% acetic acid over 83 min with a flow rate of 300 nl/min. LC-MS/MS analyses were performed on an LTQ Orbitrap Elite (ThermoFisher Scientific, Waltham, Massachusetts, USA) using an EASY-nLC 1200 liquid chromatography system. For mass spectrometry analysis, a full scan in the Orbitrap with a resolution of 60,000 was followed by collision-induced dissociation (CID) of the twenty most abundant precursor ions. MS2 experiments were acquired in the linear ion trap.

### MS data analysis

A database search against a *Salmonella* Typhimurium SL1344 annotation downloaded from Uniprot (date 23/08/2018, organism ID 216597, 4,659 entries) as well as label-free quantification (LFQ) was performed using MaxQuant (version 1.6.2.6) (Cox and Mann, 2008). Common laboratory contaminants and reversed sequences were included by MaxQuant. Search parameters were set as follows: Trypsin/P specific digestion with up to two missed cleavages, methionine oxidation (+15.99 Da) as a variable and carbamidomethylation at cysteines (+57.02 Da) as fixed modification, match between runs with default parameters enabled. The FDRs (false discovery rates) of protein and PSM (peptide spectrum match) levels were set to 0.01. Two identified peptides with at least one of those being unique were required for protein identification. LFQ was performed using the following settings: LFQ minimum ratio count 2 considering only unique for quantification. Results were filtered for proteins quantified in at least two out of three biological replicates before statistical analysis. Here, two strains (of either wild-type, Δ*mgrB*, *mgrB*+) were compared by a student’s *t*-test applying a threshold p-value of 0.01, which was based on all possible permutations.

### Infection assay, CFU counting, and flow cytometry

For infection of HeLa cells, overnight cultures were diluted 1:100 in fresh LB medium (supplemented with the appropriate antibiotic if needed) and grown aerobically to an OD_600_ of 2.0. Bacteria were collected by centrifugation (2 min at 12,000 rpm, RT) and resuspended in DMEM. HeLa cells were infected at a multiplicity of infection (MOI) of 25. After addition of the bacteria, cells were centrifuged at 250*g* for 10 min at RT and subsequently incubated at 37°C in 5% CO_2_ and humidified atmosphere. After this, the medium was exchanged (this step marking the time zero) with one containing 50 μg/ml gentamicin for 30 min to kill extracellular bacteria, and then changed again to 10 μg/ml for the remainder of the infection. Cells were collected at the indicated time points. Colony forming unit (CFU) assays were performed to quantify intracellular bacteria. At the indicated time points, the infected cells were washed twice with PBS and lysed in PBS with 0.1% (v/v) Triton X-100. The lysates were diluted in PBS and plated on LB plates (with appropriate antibiotic if required) and incubated overnight at 37°C. CFUs were then counted and normalized to the number of the bacterial inoculum used for the infection and fold-changes relative to the wild-type strain for each time point were calculated. The flow cytometry assay was used to quantify the bacterial intracellular replication. GFP-expressing *Salmonella* were used for infection of HeLa cells as described. Infected cells were detached at the indicated time points by trypsin digestion (manufacturer), washed with cold PBS by centrifugation (5 min, 550*g* at 4°C) and analysed with a FACS BD Accuri^TM^ C6 (BD Biosciences) instrument, collecting at least 20,000 GFP-positive cells for each condition. Fluorescence was then compared to 1 h-infected cells. The data were analysed with the FlowJo software (Tree Star Inc.). Infection of RAW.B macrophages was carried out as in the TraDIS protocol.

### Motility assay

For swimming assays, 6 μl of overnight cultures in LB or SPI-2 minimal medium were spotted at the centre of 0.3% agar SPI-2 plates, in the presence of the appropriate antibiotic if required. Swimming was monitored after 6 h of incubation at 37°C.

### Rifampicin assay

For the RNA stability assay, rifampicin was added at a final concentration of 500 μg/ml to liquid cultures in SPI-2 medium at an OD_600_ 0.3. Samples for total RNA were collected at the indicated time points following rifampicin addition, and then RNA was extracted and analysed by Northern blot as described above.

## DECLARATIONS

### Ethics approval

Not applicable

### Consent for publication

Not applicable

### Availability of data and materials

All sequencing and proteomics data have been uploaded on repositories and are available upon requests.

### Authors’ contributions

EV, AJW, and JV designed the research. EV, SLS, AKC performed lab work. EV, SM, RG, FE, LL, and LB analysed data. EV, SLS, AJW, and JV interpreted the data and wrote the manuscript, which all co-authors revised.

## Acknowledgements

We thank Barbara Plaschke for technical assistance.

